# Transcriptomic Landscape of Microglia in Mouse Models of Social Dysfunction and Oxytocin-Mediated Recovery

**DOI:** 10.64898/2026.02.26.708356

**Authors:** Masafumi Tsurutani, Haruaki Sato, Mitsue Hagihara, Dooseon Cho, Mitsutaka Kadota, Takefumi Kondo, Kazunari Miyamichi

## Abstract

Atypical sociability is a hallmark of neurodevelopmental disorders arising from genetic susceptibility and prenatal environmental perturbations affecting diverse brain cell types. Using single-cell transcriptomics, we previously identified selective vulnerability of parvocellular oxytocin (OT) neurons in the paraventricular hypothalamus (PVH) following embryonic exposure to valproic acid (VPA), a teratogen that induces social deficits. Neonatal chemogenetic activation of OT neurons rescued these behavioral abnormalities and partially restored dysregulated gene expression. However, the effects of VPA exposure and OT neuron stimulation on non-neuronal PVH cells remained unclear. Here, we show that VPA induces transcriptional abnormalities in PVH microglia. Spatial transcriptomics revealed altered distributions of PVH microglial subtypes. Notably, neonatal OT neuron stimulation reversed a subset of VPA-induced microglial gene downregulation, while pharmacological manipulation of microglia normalized aberrant *OT* gene expression in putative parvocellular OT neurons. These findings support bidirectional OT neuron–microglia interactions that may underlie social dysfunction following embryonic VPA exposure.

**Highlights:**

- Embryonic VPA exposure induces potent transcriptional changes in PVH microglia
- PVH microglia comprise two spatially organized subtypes disrupted by VPA
- Neonatal OT neuron stimulation restores gene expression in a microglial subtype
- Microglial manipulation rescues OT ligand in parvocellular PVH neurons

## Introduction

Non-neuronal cells constitute a substantial fraction of brain tissue and play essential roles in neural homeostasis^1,2^; their dysfunction is increasingly recognized as a contributor to brain disorders. Microglia, the resident immune cells of the central nervous system, are critical for neurodevelopmental processes and for lifelong maintenance of neural circuits^3^. Accordingly, microglia have been implicated in neurodevelopmental disorders, including autism spectrum disorder (ASD)^3^, as well as neurodegenerative diseases such as Alzheimer’s and Parkinson’s disease^4^. Although microglia were traditionally conceptualized within an M1 (pro-inflammatory/neurotoxic) versus M2 (anti-inflammatory/neuroprotective) dichotomy^5^, recent single-cell transcriptomic studies have revealed greater heterogeneity and region-dependent specialization among microglial populations^4^. These advances underscore the need for systematic characterization of microglial states across brain regions and pathological conditions. In this study, we provide a focused analysis of microglia in the paraventricular hypothalamus (PVH), in relation to social behavior and the oxytocin (OT) system^6^.

OT is a nonapeptide hormone produced by OT neurons located in the PVH and supraoptic hypothalamus. OT released from the PVH is involved in social behaviors, bond formation, and parental behaviors^6–9^. Within the PVH, magnocellular OT neurons predominantly project to the posterior pituitary, whereas parvocellular OT neurons send selective projections to central brain regions^10,11^. Transcriptomic analyses distinguish these neuronal subtypes^10,12^, with parvocellular OT neurons thought to play a dominant role in regulating social behaviors^10^. Impaired OT neuron function has been documented in genetic and environmental rodent models of social dysfunction, including *Shank3b* mutants and prenatal valproic acid (VPA) exposure^12–14^. In our previous study, single-nucleus RNA sequencing (snRNAseq) of the PVH in mice exposed to VPA during embryogenesis revealed that parvocellular PVH OT neurons exhibit dysregulation of key signal-transduction pathways and reduced *OT* gene expression, whereas magnocellular PVH OT neurons were comparatively spared^12^. Notably, chemogenetic stimulation of OT neurons during the neonatal or young-adult period ameliorated social deficits and restored *OT* expression in the VPA-exposed model^12^. Despite evidence that microglia contribute to the regulation of social behaviors^15–18^, it remains unclear how embryonic VPA exposure affects non-neuronal cell types in the PVH. These considerations motivated reanalysis of the snRNAseq dataset and incorporation of spatial transcriptomics with a focus on non-neuronal cell populations.

## Results

### Impact of embryonic VPA exposure on non-neuronal PVH cells

To assess the effects on non-neuronal cell populations in a mouse model of social deficits induced by VPA exposure on embryonic day 12.5^13,19^, we reanalyzed a previously generated snRNAseq dataset (10X Genomics Chromium) comprising 10,060 nuclei from VPA-treated mice and 3,750 nuclei from saline controls (Figure 1A)^12^. Based on canonical marker genes, PVH cells were classified as neurons, astrocytes, oligodendrocytes, oligodendrocyte precursor cells (OPCs), microglia, and ependymal cells (Figure S1A, B). Differentially expressed genes (DEGs) were defined by an absolute log_2_ fold change (|log2-FC|) > 0.5 and a p < 0.05. Both up- and downregulated DEGs were more numerous in astrocytes, oligodendrocytes/OPCs, and microglia than in neurons, indicating broad transcriptomic effects of embryonic VPA exposure on non-neuronal cells (Figure S1C, D).

**Figure 1:**
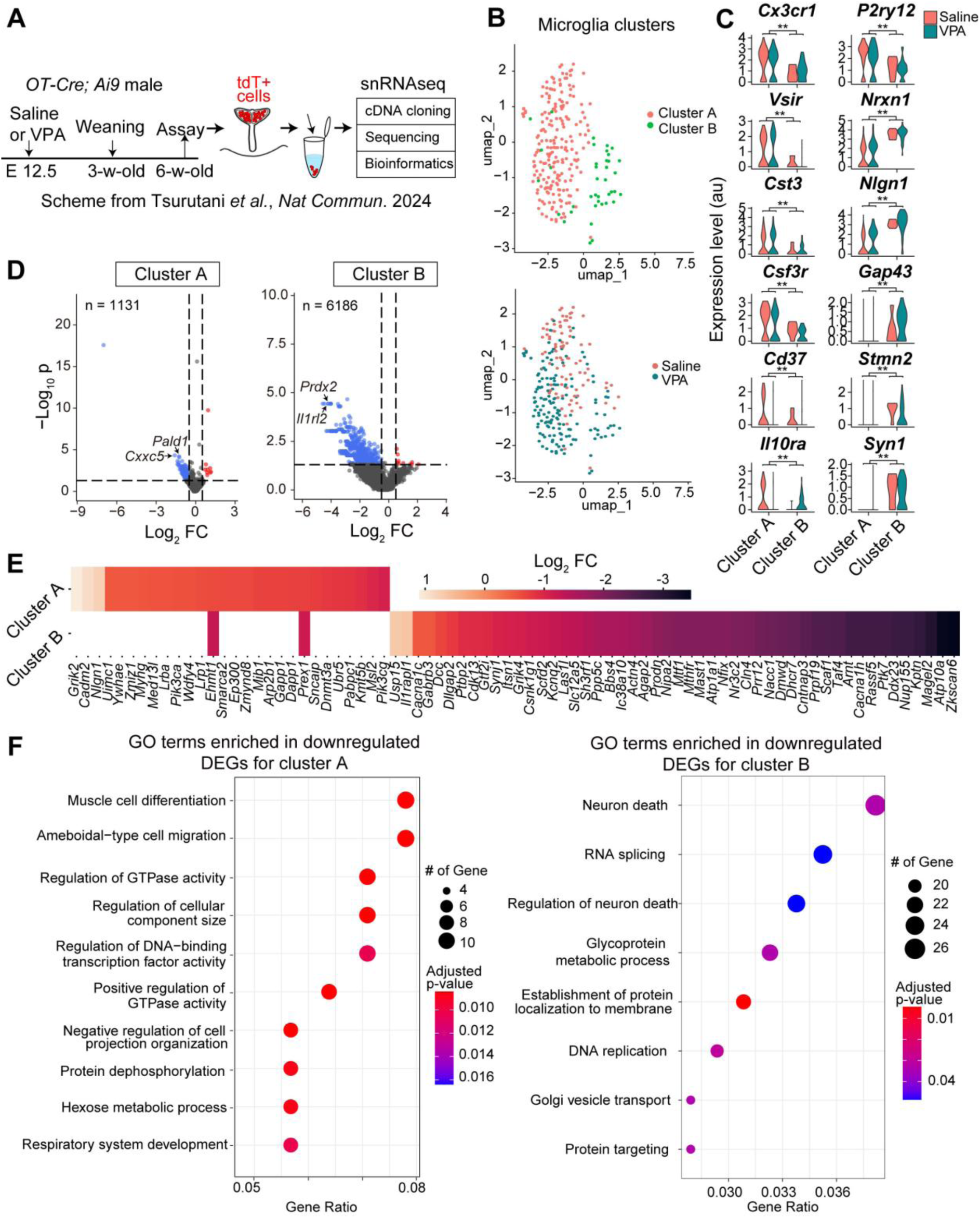
Transcriptomic profiles of PVH microglia in VPA-treated mice. (A) Schematic of snRNAseq data collection and analysis, as described previously^12^. tdT, tdTomato from the *Ai9* allele. (B) Top: UMAP representation of PVH microglia showing two clusters (A and B). Bottom: Both saline- and VPA-treated groups contained these two microglial clusters. (C) Violin plots showing representative marker genes distinguishing microglial clusters A and A. **, p < 0.01 by the Wilcoxon rank-sum test. Of note, the expression levels of these genes did not differ significantly between the saline- and VPA-treated groups (p > 0.05 by the Wilcoxon rank-sum test). (D) Volcano plots showing DEGs within each cluster (upregulated in the VPA-treated group, red; downregulated in the VPA-treated group, blue). The *x*-axis indicates log_2_ fold change, and the *y*-axis indicates −log_10_ p-value. Numbers of genes analyzed (n) are shown. P-values were calculated using the Wilcoxon rank-sum test without adjustment for multiple comparisons. (E) Heatmaps showing log_2_ fold changes of ASD risk factor DEGs identified in PVH microglia. (F) Heatmap of GO terms (p < 0.05) enriched among downregulated DEGs in microglial clusters A and B. P-values were calculated using a one-sided Fisher’s exact test with Benjamini–Hochberg correction. See Figure S1 for additional data.

Given evidence that microglia regulate social behaviors^15–18^, we focused on PVH microglia. Uniform Manifold Approximation and Projection (UMAP) revealed two transcriptomic microglial clusters (hereafter referred to as clusters A and B), each containing cells from saline- and VPA-treated mice (Figure 1B) and expressing canonical microglial markers, including *CX3C motif chemokine receptor 1* (*Cx3cr1*) and *purinergic receptor P 2y12* (Figure 1C). Marker gene analysis showed that cluster A exhibited elevated *Cx3cr1* and *P 2y12*, whereas cluster B expressed higher levels of neuronal-associated genes, including *Neurexin 1* (*Nrxn1*) and *Synapsin 1* (*Syn1*) (Figure 1C). P2RY12 expression is thought to be elevated in activated microglia^20^. Consistent with this distinction, gene ontology (GO)^21,22^ and pathway^23^ analyses indicated immune-related functions for cluster A and synaptic or neuron-associated signatures for cluster B (Figure S1E, F). Transcription factor activity analysis^24^ revealed distinct regulatory profiles between the PVH microglial subtypes (Figure S1G–I). In both clusters, VPA-treated mice showed more downregulated than upregulated DEGs (Figure 1D), indicating a predominantly suppressive transcriptomic effect of embryonic VPA exposure on microglia.

We next examined DEGs previously designated as high-confidence ASD risk genes^25^ and identified distinct sets in microglial clusters A and B, with minimal overlap (Figure 1E). GO analyses of downregulated DEGs by prenatal VPA exposure indicated enrichment for GTPase activity and cell-size regulation in cluster A, whereas cluster B was enriched for neuronal cell death and RNA splicing terms (Figure 1F). Together, these findings indicate that the two PVH microglial subtypes are differentially affected by embryonic VPA exposure.

### Spatial transcriptomic profiling of microglial subtypes within and near the PVH

Recent advances in spatial transcriptomics (STx) have begun to reveal the spatial organization of transcriptomic subtypes across the brain^26–28^. We reasoned that these approaches would enable detailed characterization of microglial subtype organization within the PVH and reveal how embryonic VPA exposure perturbs these spatial patterns. To address these points, we performed STx profiling of PVH cells using the 10X Xenium platform^29^ on sections from saline- or VPA-treated mice. Eighteen sections from nine mice (N = 4 and 5 for the saline and VPA-treated groups, respectively) were processed with the Xenium Mouse Gene Expression Panel (Figure S2A). After quality control and cell segmentation, 175,755 cells within and adjacent to the PVH were identified and classified into 46 transcriptomic clusters (Tables S1, S2). Based on canonical marker genes, cells were annotated as neurons (14 PVH subtypes and nine additional subtypes elsewhere), astrocytes (14 subtypes), oligodendrocytes/OPCs (five subtypes), microglia (two subtypes), and ependymal cells (two subtypes) (Figure S2B–D). Spatial distributions of PVH neuronal subtypes are shown in Figure S3.

From this dataset, we identified 5,153 microglia (2.9% of detected cells) within and adjacent to the PVH (Figure 2A). Based on representative marker genes (Figure 2B), STx subtype #20, characterized by elevated *Syn1* expression, corresponded to neuronal-type microglia (snRNAseq cluster B), whereas STx subtype #31, showing high *Cx3cr1* expression, corresponded to immune-type microglia (snRNAseq cluster A). In saline-treated mice, immune-type microglia (subtype #31) were enriched in the anterior PVH, where magnocellular OT neurons predominate^10,12^, whereas neuronal-type microglia (subtype #20) were more frequent in the posterior PVH, which contains a higher proportion of parvocellular OT neurons^10,12^. This spatial distribution was reversed in VPA-exposed mice, with increased subtype #20 abundance in the anterior PVH and predominance of subtype #31 in the posterior PVH (Figure 2C).

**Figure 2:**
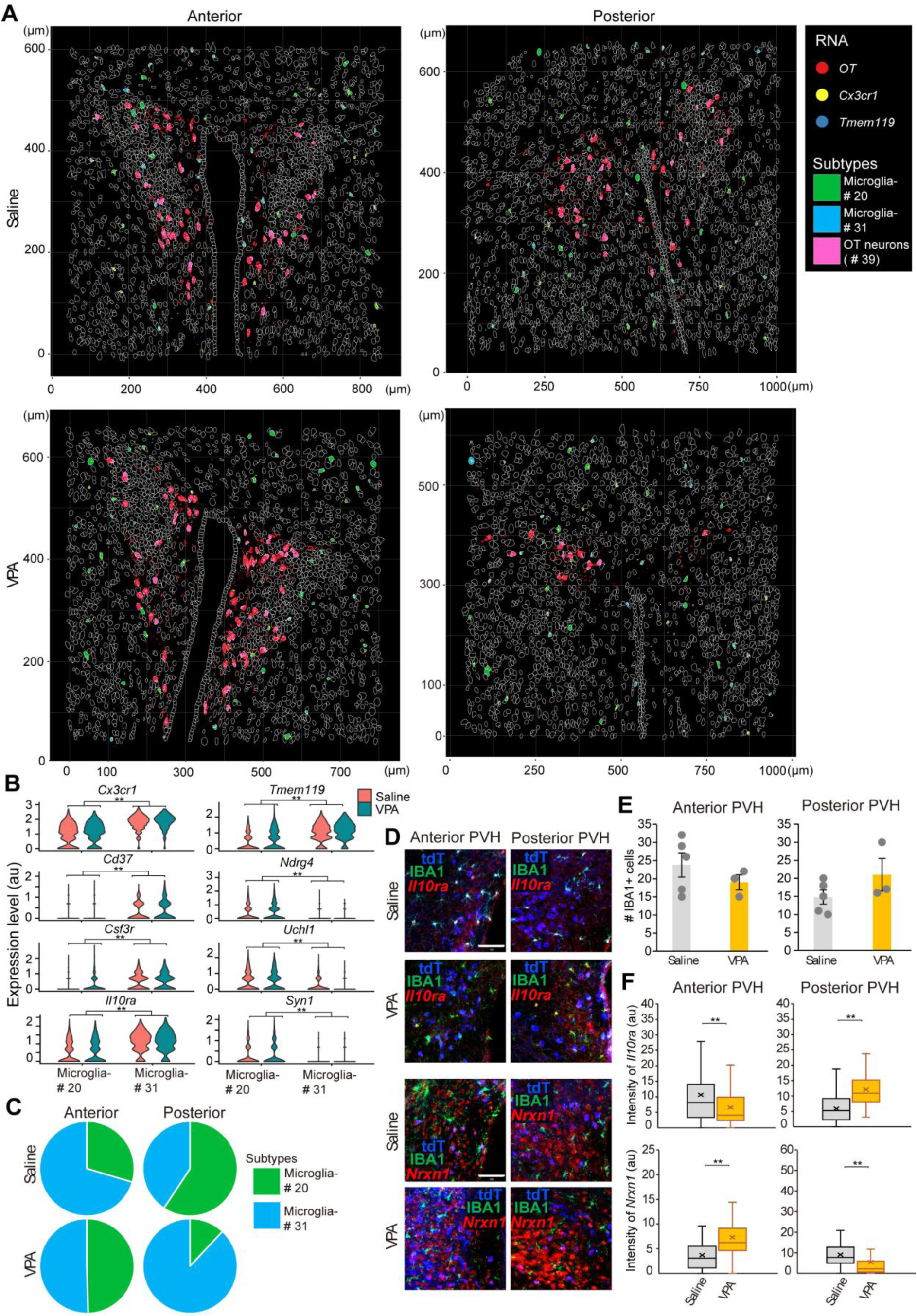
Spatial transcriptomic analysis of microglial subtypes in the PVH. (A) Spatial map of OT neurons (pink) and two microglial subtypes (green and blue) identified using the 10X Xenium platform. White contours delineate detected cells in the anterior and posterior PVH. Dots indicate individual RNA signals (red, *OT*; yellow, *Cx3cr1*; indigo, *Tmem119*). (B) Violin plots showing representative marker genes distinguishing microglial subtypes. Subtype #20 corresponds to neuronal-type microglia (snRNAseq cluster B), whereas subtype #31 corresponds to immune-type microglia (snRNAseq cluster A). **, p < 0.01 by the Wilcoxon rank-sum test. (C) Pie charts showing proportions of the two microglial subtypes in the anterior and posterior PVH of saline- and VPA-treated mice. (D) Representative coronal PVH sections from saline- and VPA-treated mice showing IBA1 immunostaining and expression of the indicated marker genes (*Il10ra* and *Nrxn1*) detected by *in situ* hybridization (ISH). Scale bar, 100 μm. (E) Quantification of IBA1-positive cell numbers in the PVH. No significant difference was detected by the two-sided Wilcoxon rank-sum test (N = 5 saline, N = 3 VPA). (F) Fluorescence intensity of ISH signals for *Il10ra* and *Nrxn1*. A total of 59–121 cells from three animals per condition were analyzed. **, p < 0.01 by the two-sided Wilcoxon rank-sum test. Box plots show the median (center line), first and third quartiles (box boundaries), and whiskers extending to 1.5× the interquartile range. See Figures S2 and S3 for additional data on other cell types.

To validate these observations in an independent cohort, we performed immunostaining for IBA1, a canonical microglial marker^30^, together with *in situ* hybridization (ISH) for *Il10ra* (a marker of subtype #31) and *Nrxn1* (a marker of subtype #20) (Figure 2D). The number of IBA1-positive microglia did not differ between saline- and VPA-treated mice (Figure 2E). However, *Il10ra* expression was enriched in the anterior PVH of saline controls but shifted to the posterior PVH in VPA-treated mice (Figure 2F). Conversely, *Nrxn1* expression, typically higher in the posterior PVH of control mice, became more prominent in the anterior PVH following VPA exposure (Figure 2F). These histochemical findings are consistent with the STx data and support the conclusion that embryonic VPA exposure disrupts the spatial organization of microglial subtypes within the PVH.

### Impact of neonatal OT neuron stimulation on transcriptomic profiles of non-neuronal cells

Our previous work demonstrated that a single chemogenetic stimulation of OT neurons at postnatal day (PND) 2 rescues social deficits and aberrant *OT* gene expression in VPA-treated mice^12^. To examine how neonatal OT neuron stimulation affects non-neuronal populations, we reanalyzed snRNAseq datasets from VPA-treated mice with or without neonatal OT neuron stimulation (Figure 3A). Neonatal clozapine-N-oxide (CNO) treatment of VPA-treated *OT-Cre*; *stop-hM3* mice induced more upregulated than downregulated DEGs across neurons, astrocytes, and microglia, with the strongest upregulation observed in microglia (Figure S4A, B). When microglia were subdivided into clusters A and B, both saline- and CNO-treated groups contained these clusters (Figure 3B, C). In both clusters, CNO treatment resulted in more upregulated than downregulated DEGs (Figure 3D). GO analysis indicated enrichment of synapse-organization terms in cluster A, whereas neuron-death-related terms predominated in cluster B, suggesting differential modulation of microglial subtypes by neonatal OT neuron activation. In an independent cohort with ISH for *Il10ra* mRNA expression, a cluster A marker, in IBA1-expressing microglia was reduced in the posterior PVH and showed a trend toward increased expression in the anterior PVH following CNO treatment (Figure S4C, D), consistent with partial restoration of the anterior–posterior distribution of cluster A microglia in VPA-exposed mice.

**Figure 3:**
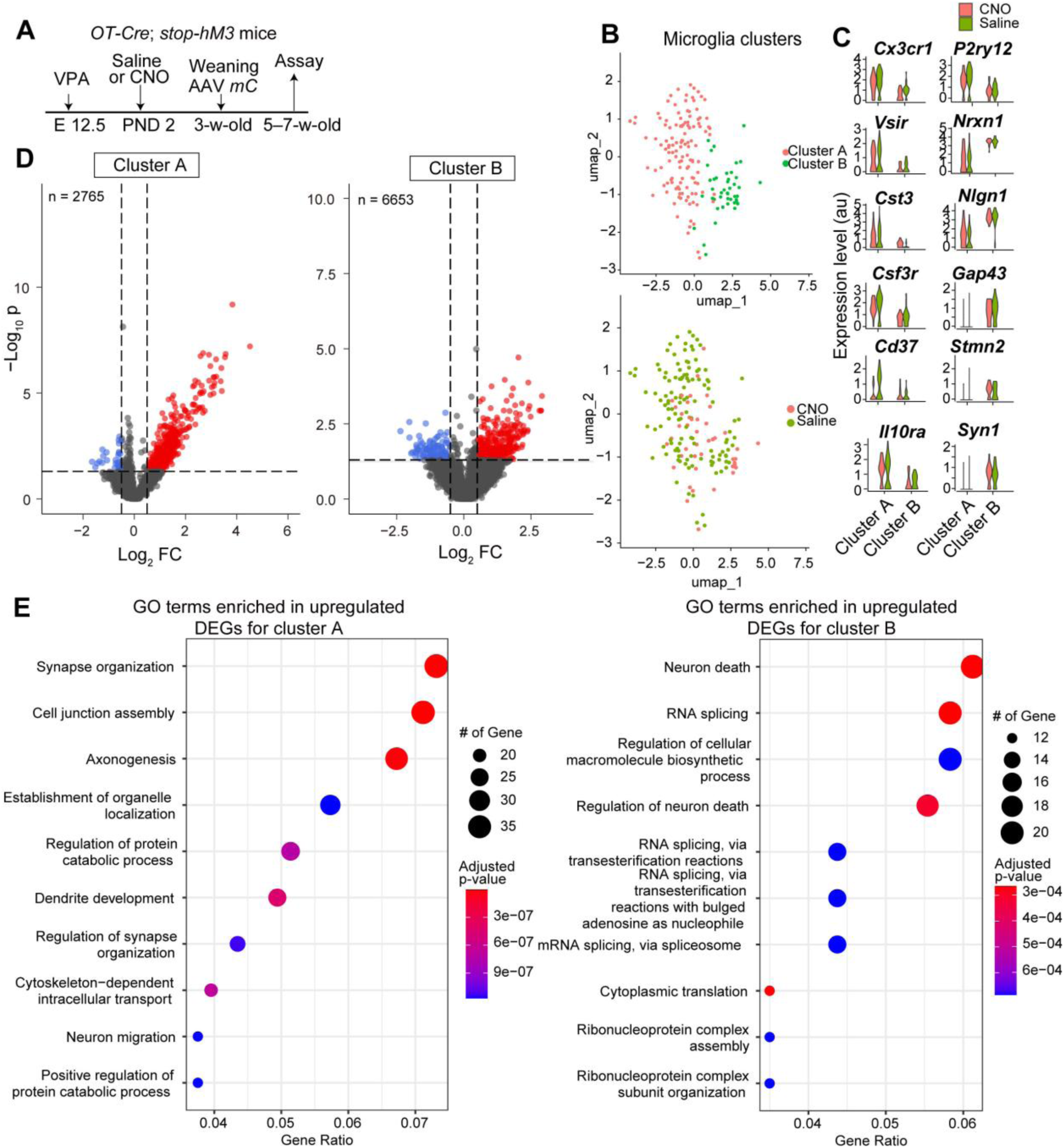
Neonatal OT neuron stimulation modulates PVH microglial transcriptomic signatures in VPA-treated mice. (A) Experimental timeline, as described previously. PND, postnatal day; mC, mCherry. (B) Top: UMAP representation of PVH microglia showing two clusters (A and B). Bottom: Both saline- and CNO-treated groups contained these two microglial clusters. (C) Violin plots showing representative marker genes distinguishing microglial clusters A and B. (D) Volcano plots showing DEGs within each cluster (upregulated in the CNO-treated group, red; downregulated in the CNO-treated group, blue). The *x*-axis indicates log_2_ fold change and the *y*-axis indicates −log_10_ p-value. Numbers of genes analyzed (n) are shown. P-values were calculated using the Wilcoxon rank-sum test without adjustment for multiple comparisons. (E) Heatmap of GO terms (p < 0.05) enriched among upregulated DEGs in microglial clusters A and B. P-values were calculated using one-sided Fisher’s exact test with Benjamini–Hochberg correction. See Figure S4 for additional data.

Given these CNO-induced transcriptomic changes, we compared gene expression alterations driven by embryonic VPA exposure and neonatal CNO treatment. Scatter plots in Figure 4A depict the relationship between prenatal VPA-induced (*y*-axis) and neonatal CNO-induced (*x*-axis) fold changes. Cluster B contained more DEGs overall. A substantial subset of genes was downregulated by VPA and upregulated by CNO (red dots in Figure 4A), indicating reversal of VPA-induced abnormalities; representative examples are shown in Figure 4B. GO enrichment analysis revealed that reversed DEGs in cluster B were associated with neural cell death and RNA regulation, whereas no significant GO terms were enriched in cluster A (Figure 4C). Conversely, genes upregulated by VPA and downregulated by CNO were infrequently observed (green dots in Figure 4A). These findings indicate that embryonic VPA exposure induces transcriptional downregulation and that neonatal chemogenetic activation of OT neurons partially restores these changes, predominantly in microglial cluster B.

**Figure 4:**
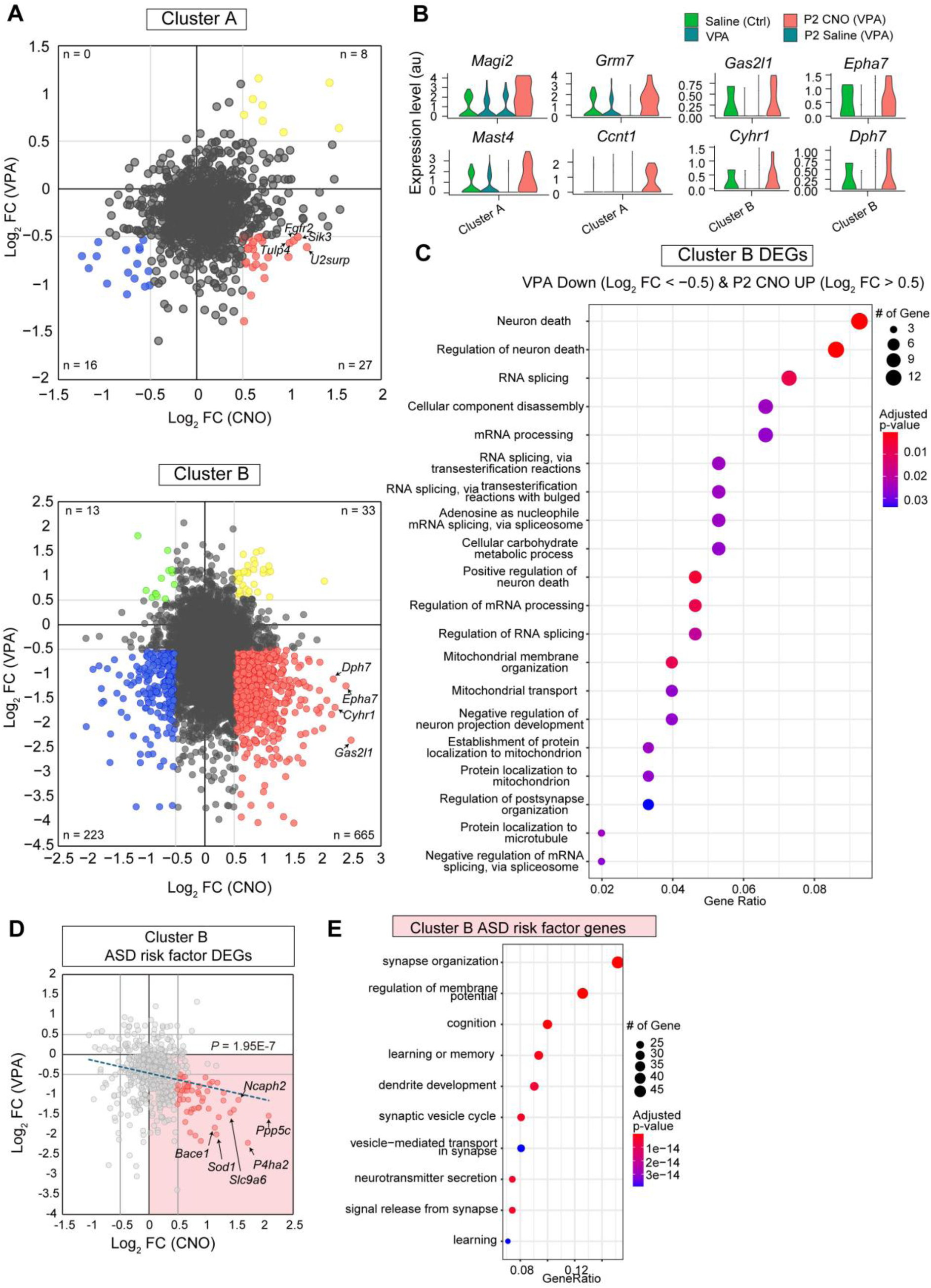
Integrated analysis of PVH microglial transcriptomic signatures in social dysfunction and OT-mediated recovery models. (A) Scatter plots showing correlations between log_2_ fold changes in microglial gene expression induced by embryonic VPA exposure (*y*-axis) and neonatal CNO treatment (*x*-axis) for microglial cluster A (top) and cluster B (bottom). Red dots indicate DEGs downregulated by VPA and upregulated by CNO; green dots indicate DEGs upregulated by VPA and downregulated by CNO; yellow and blue dots represent DEGs up- and downregulated under both conditions, respectively. DEGs were more enriched in cluster B. (B) Violin plots showing representative cluster-distinguishing marker genes corresponding to red-dot DEGs in panel A. (C) Heatmap of GO terms (p < 0.05) enriched among red-dot DEGs in cluster B. P-values were calculated using one-sided Fisher’s exact test with Benjamini–Hochberg correction. (D) Scatter plots showing correlations between log_2_ fold changes in ASD risk factor genes induced by embryonic VPA exposure (*y*-axis) and neonatal CNO treatment (*x*-axis) for microglial cluster B. Light red dots (49 genes) indicate DEGs downregulated by VPA and upregulated by CNO with some gene names indicated. The p-value is based on one-sided t-test. (E) Heatmap of GO terms (p < 0.05) enriched among genes located in the light red area of panel D (including 329 genes). The p-value was calculated using one-sided Fisher’s exact test with Benjamini–Hochberg correction. See Figures S5 and S6 for additional data.

To further assess the potential functional restoration of microglial cluster B, we analyzed ASD risk factor genes expressed in this cluster. Overall, we observed a significant trend in which ASD risk factor genes that were downregulated by prenatal VPA exposure were upregulated following neonatal CNO treatment (Figure 4D). Notably, GO analysis of significantly restored 329 gene sets (corresponding to red dots in Figure 4D) demonstrated enrichment for synaptic and neuron-related categories (Figure 4E). In addition, NeuronChat^31^ analysis suggested potential OT neuron–microglia interactions involving cluster B via neurexin–neuroligin signaling, with no detectable interactions between OT neurons and microglial cluster A (Figure S5). Consistent with the absence of predicted OT–OT receptor (OTR) interactions, neither STx nor histochemical analyses showed evidence for widespread OTR expression in microglia (Figure S6). Together, these data suggest a potential functional restoration of microglial cluster B following neonatal OT neuron stimulation, possibly mediated by their molecular interactions between OT neurons and microglia, although indirect mechanisms cannot be excluded (see Discussion).

### Impacts of microglial manipulations on OT ligand expression

We next examined whether experimental manipulation of microglia ameliorates OT ligand downregulation caused by embryonic VPA exposure. Two pharmacological approaches were used. First, minocycline, a tetracycline-class antibiotic that inhibits microglial activation^32^, has been reported to rescue VPA-induced sociability deficits in rats^33^. Second, the colony-stimulating factor 1 receptor (CSF1R) antagonist PLX5622, which depletes resident microglia followed by repopulation with naïve microglia, improves microglial function^34^. Based on these findings, VPA-exposed mice were treated with minocycline from 4 weeks of age for two weeks, followed by behavioral assessment (Figure 5A, B). Whereas vehicle-treated mice exhibited reduced social interaction in the three-chamber test, minocycline treatment improved sociability, consistent with previous reports^33^. Similarly, PLX5622 treatment from 4 weeks of age for two weeks, followed by a one-week recovery period, restored sociability in VPA-exposed mice (Figure 5B). Although these experiments do not identify the specific microglial subtypes or anatomical loci involved, the results are consistent with prior evidence that microglial manipulation exerts pro-social effects.

**Figure 5:**
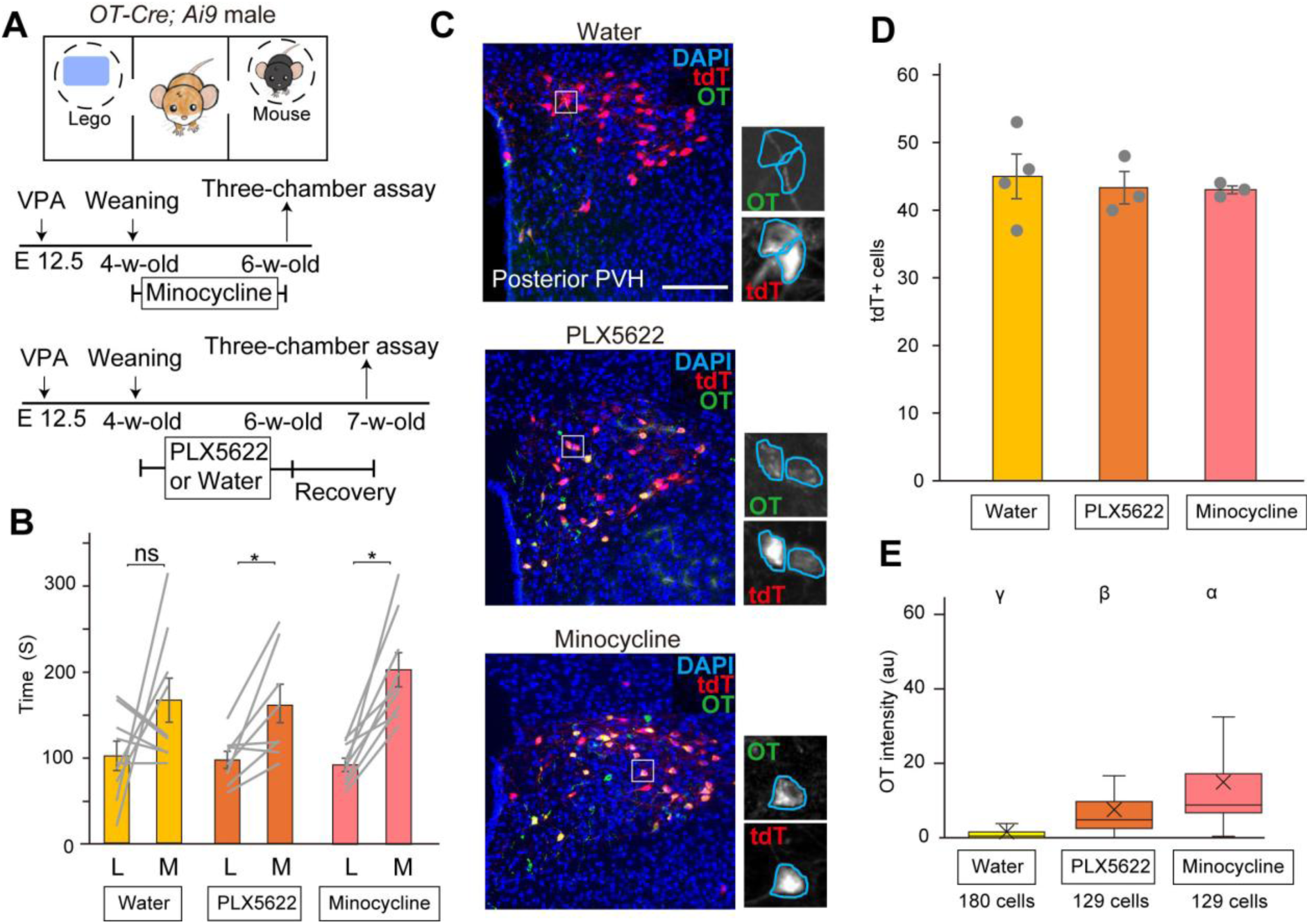
Long-lasting effects of microglia manipulation on sociability and OT expression. (A) Experimental timeline for microglial manipulations. (B) Time spent exploring a Lego block (L) or an unfamiliar male mouse (M) in the three-chamber sociability assay. N = 9 for water control and minocycline-treated groups and N = 8 for the PLX5622-treated group. *p < 0.05 by the two-sided Wilcoxon rank-sum test. (C) Representative coronal PVH sections showing tdTomato-positive cells (tdT, red) and anti-OT immunostaining (OT, green), counterstained with DAPI. Scale bars, 200 μm (low magnification) and 20 μm (high magnification). (D) Number of tdT-positive cells per PVH section. N = 4 for the water group and N = 3 for PLX5622- and minocycline-treated groups. (E) Fluorescence intensity of anti-OT staining. The indicated numbers of cells were quantified from N = 3 animals per group. Box plots are as in Figure 2F. Different letters (α, β, γ) indicate p < 0.05 by the two-sided Wilcoxon rank-sum test with Bonferroni correction. Box plots follow the definitions in Figure 2F. Error bars indicate SEM. See Figures S7–S9 for additional data.

Because VPA exposure also produces non-social behavioral abnormalities, we examined repetitive, compulsive-like behaviors using the marble-burying test^35^ (Figure S7A). As reported previously, VPA-treated mice buried more than 10 marbles, compared with approximately three to four marbles buried by controls^12^. In the present cohort, vehicle-treated VPA mice buried more than 10 marbles, whereas minocycline treatment significantly reduced marble burying (Figure S7B). In contrast, PLX5622 treatment showed variable effects and did not significantly improve repetitive behaviors. These results suggest that minocycline and PLX5622 differentially affect non-social behavioral phenotypes in VPA-treated mice.

We next assessed whether microglial manipulations modulate *OT* gene expression. Focusing on the posterior PVH, where parvocellular OT neurons reside, neither treatment altered OT neuron number, whereas both significantly increased *OT* mRNA levels (Figure 5C–E). We also examined effects on microglial gene expression and distribution. Minocycline, but not PLX5622, increased expression of *Epha7* (Figure S8A, B), a gene downregulated by VPA and restored by neonatal OT neuron stimulation (Figure 4B). *Il10ra* (a cluster A microglial marker) was upregulated in the anterior PVH following minocycline treatment but downregulated in the posterior PVH following PLX5622 treatment (Figure S8C, D). Thus, although both treatments tended to normalize aberrant gene expression in OT neurons and microglia, the magnitude and direction of individual gene changes differed.

In summary, pharmacological manipulation of microglia restored *OT* gene expression in putative parvocellular OT neurons and modulated microglial gene expression and spatial distribution, although the latter requires further validation.

## Discussion

The present study demonstrates bidirectional interactions between OT neurons and microglia in a VPA-induced model of social dysfunction (Figure S9). We identified two transcriptomic microglial subtypes in the PVH whose molecular states were differentially affected by embryonic VPA exposure and partially normalized by neonatal OT neuron stimulation, particularly within cluster B. Conversely, pharmacological manipulation of microglia upregulated OT gene expression in putative parvocellular PVH OT neurons. These reciprocal influences may contribute to the recovery of sociability and broader restoration of PVH circuit function. Below, we discuss the biological implications and limitations of this work.

Neurodevelopmental disorders characterized by atypical social behavior, including ASD, have traditionally been studied through circuit features associated with specific symptoms, particularly within the prefrontal cortex and related higher-order cortical regions^36–38^. In contrast, hypothalamic regulators of social behavior have received comparatively less attention^10,39^. Our previous study^12^ and the present work together examine transcriptomic alterations across neuronal and non-neuronal PVH cell types in both the VPA-induced social deficit model and the OT-mediated recovery model. Among these populations, microglia were particularly notable: embryonic VPA exposure induced extensive gene downregulation, much of which was reversed by neonatal OT neuron stimulation (Figure 4). These findings suggest that microglia may represent a cellular node linking parvocellular PVH OT neuron dysregulation to impaired social behavior.

Analyses of snRNAseq and STx datasets revealed two microglial transcriptomic states within the PVH. Cluster A was enriched for immune-related transcripts, including *Cx3cr1* and *Il10ra*, whereas cluster B showed higher expression of synaptic and neuron-related genes, such as *Nrxn1*, *Nlgn1*, and *Syn1* (Figures 1 and 2). Notably, neither subtype corresponded closely to previously described microglial classes in other brain regions^5,40^, suggesting the presence of PVH-specific microglial states. STx further revealed that cluster A predominated in the anterior PVH, whereas cluster B was enriched in posterior regions. These findings highlight the utility of the Xenium 5K mouse panel for resolving the spatial organization of microglial subtypes, an approach broadly applicable to mapping microglial heterogeneity across healthy, injured, and diseased brain states^40^.

Recent studies suggest that PVH microglia indirectly regulate the activity of magnocellular neurons by phagocytosing astrocytic processes that occupy the interstitial space between neighboring neurons^41^. In our data, an immune-type microglial subtype (cluster A) was preferentially distributed in the anterior PVH, where magnocellular neurons are abundant^10,12^. Thus, cluster A may represent a microglial subpopulation that modulates neuronal activity through phagocytic interactions. Because neonatal stimulation of OT neurons restored the expression of neuron-related genes in cluster B (Figure 4D, E), we speculate that cluster B may also play a role in regulating local neural circuits predominantly in the posterior PVH. In VPA-exposed mice, these anterior–posterior distribution patterns were disrupted, although the functional significance of this alteration remains unclear. Elucidating the distinct roles of each cluster will require the development of subtype-specific genetic tools to selectively manipulate PVH microglia and determine how they influence local neuronal circuits and social behavior.

How bidirectional interactions between microglia and PVH OT neurons arise remains unclear. Here, we outline several possibilities. First, because microglia are generated as early as embryonic day 9.5^42^, they may be directly affected by VPA exposure at embryonic day 12.5, either through cell-autonomous effects or activation triggered by local tissue perturbation. Aberrant microglial activation or spatial organization could secondarily impair parvocellular OT neurons, contributing to the selective deficits observed in this population. Consistent with this view, *OT* expression remains normal at PND 2 in VPA-exposed mice^12^, suggesting a delayed onset of deficits during neonatal development. Second, OT neurons may exert a restorative influence on microglia through direct molecular interactions, potentially involving Nrxn–Nlgn signaling and other ligand–receptor pairs identified by snRNAseq-based communication analysis (Figure S5). Alternatively, OT neuron activation may modulate microglia indirectly through changes in neuronal activity or local neuromodulatory tone. Third, microglial abnormalities within the PVH may represent a consequence rather than a cause of impaired social behavior. For example, atypical social interactions with the dam or littermates in VPA-exposed pups could impose early-life social stress that secondarily alters PVH microglia. Distinguishing among these possibilities and establishing causality will require cell-type- and region-specific perturbations of microglia with higher temporal precision.

We acknowledge some limitations of the present study. First, the two transcriptomic subtypes of PVH microglia were defined using snRNAseq, STx, and a limited set of histochemical markers, without establishing links to cellular morphology or cytological features using *ex vivo* approaches. Second, we examined only the VPA-induced social deficit model, leaving unresolved whether the microglial alterations identified here generalize to genetic ASD models, such as *Shank3b* mutants. Third, STx was used exclusively to characterize VPA-induced abnormalities; therefore, the effects of neonatal OT neuron stimulation on the spatial organization of PVH cells, including anterior–posterior patterning of microglial subtypes, remain uncertain, although partial restoration was inferred from a subset of marker genes (Figure S4C, D). Similarly, the impact of microglial manipulations was assessed only by histochemistry, and analyses using snRNAseq or STx remain to be performed. Accordingly, the STx analysis presented here should be regarded as preliminary. Nevertheless, our combined snRNAseq- and STx-based analyses of non-neuronal PVH cells provide a foundation for future studies aimed at comprehensively mapping transcriptomic and spatial alterations across diverse mouse models of social deficits.

## STAR★Methods

### Key resources table

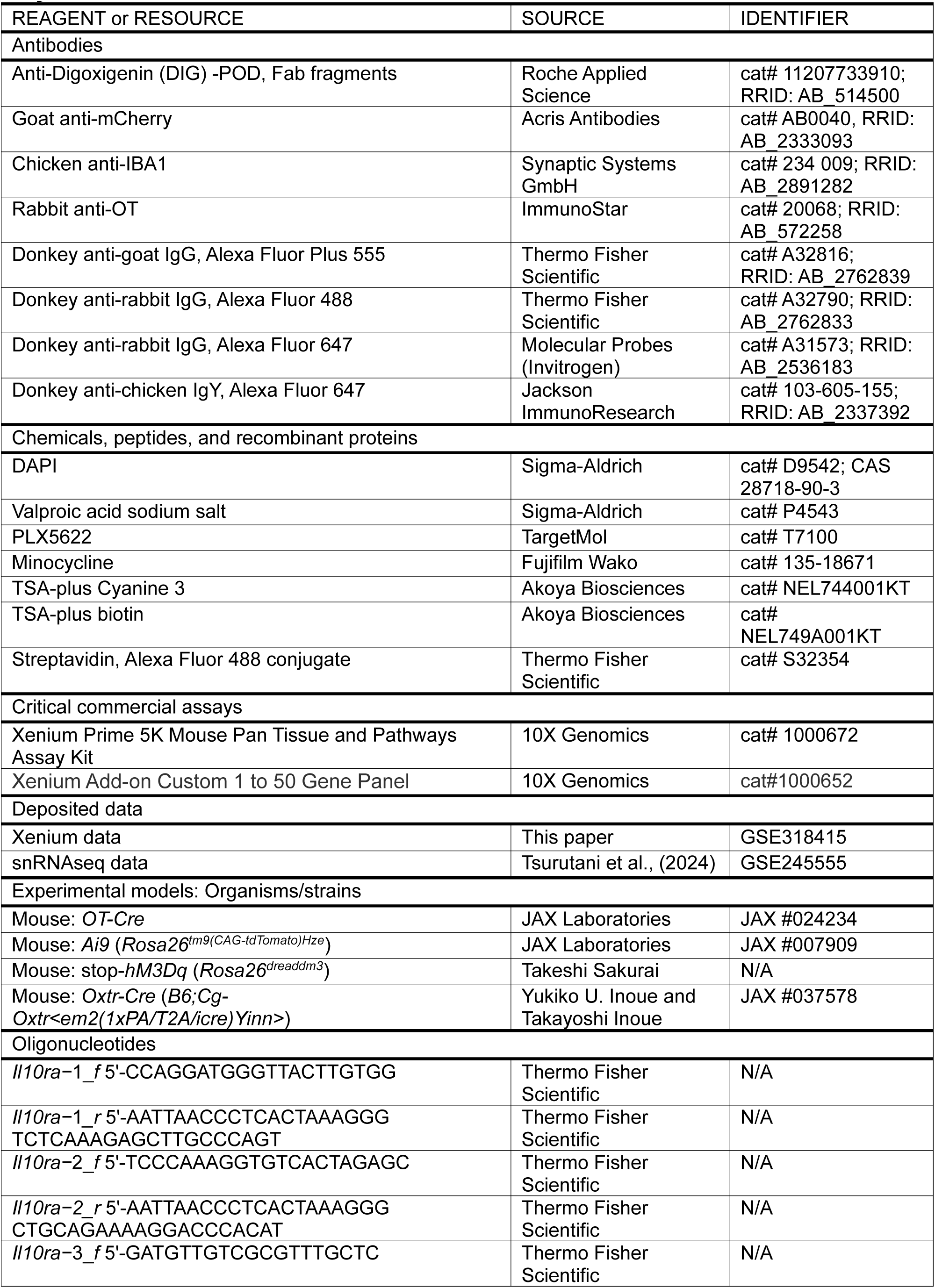
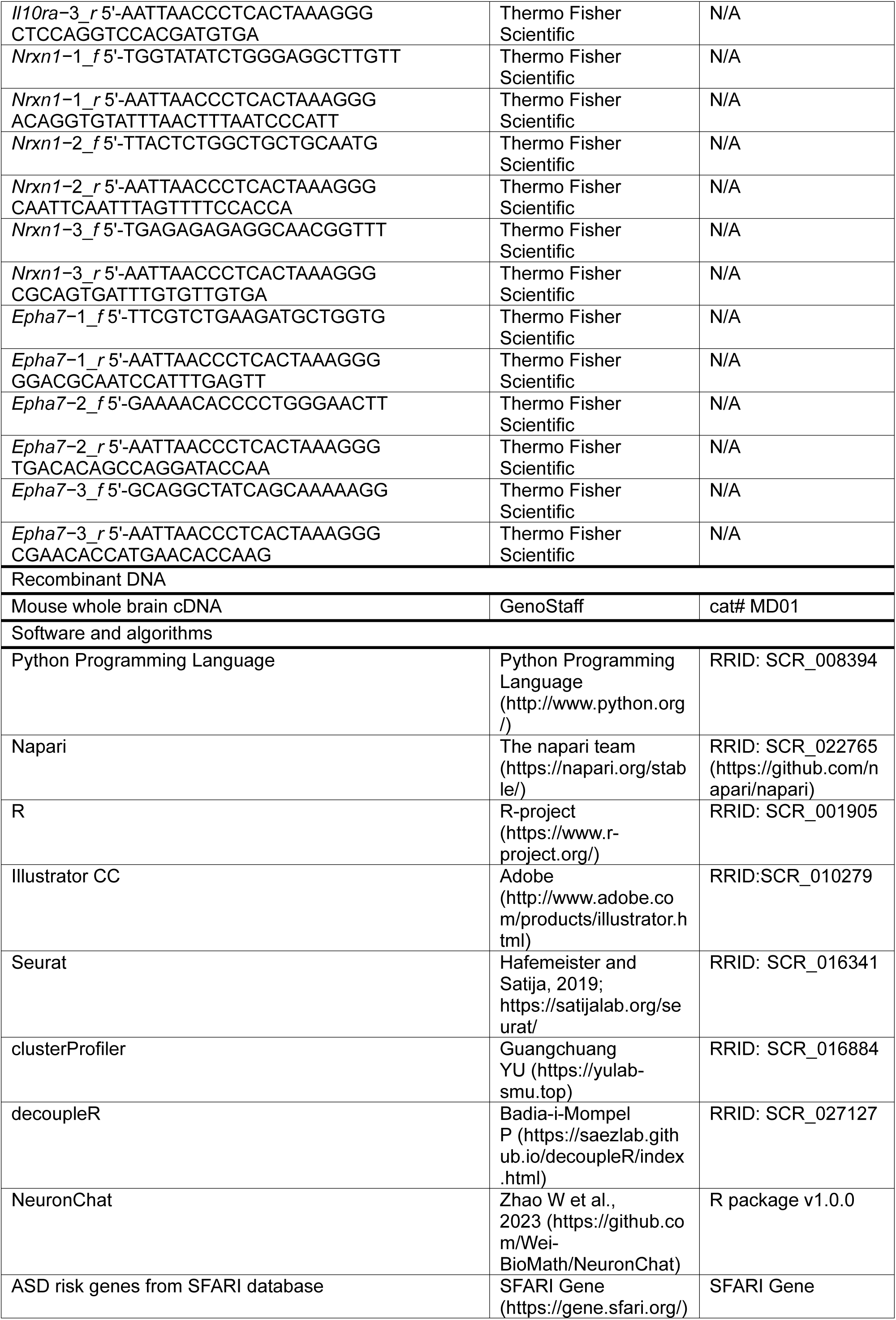
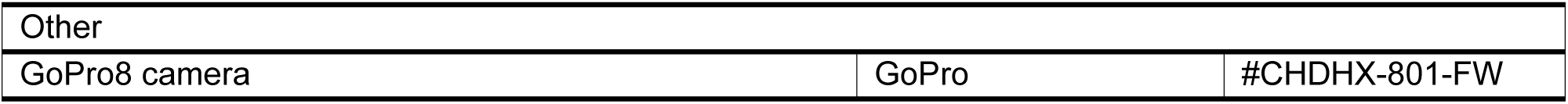

### Experimental Model and Subject Details

#### Animals

All animal procedures were approved by the Institutional Animal Care and Use Committee of the RIKEN Kobe Branch (protocol #A2017-15-13). *OT-Cre* mice (JAX #024234) were purchased from The Jackson Laboratory. *Rosa26^tm9(CAG-tdTomato)Hze^* (*Ai9*) reporter mice (JAX #007909) were provided by Dr. Takeshi Imai, who originally obtained them from the Jackson Laboratory. *Rosa26^dreaddm3^* mice (also known as stop-*hM3Dq* mice) were provided by Takeshi Sakurai. *Oxtr-Cre* (RIKEN BRC11687, also known as B6; C3-*Oxtr^em2(icre)Yinn^*/J, Jax #037578) mice were provided by Dr. Yukiko U. Inoue and Dr. Takayoshi Inoue. Wild-type C57BL/6J females used for breeding were purchased from Japan SLC, Inc. (Shizuoka, Japan). All mouse lines were maintained on a C57BL/6J background. Animals were housed at the RIKEN Center for Biosystems Dynamics Research (BDR) under standard conditions (ambient temperature 18–23 °C; relative humidity approximately 45%) with a 12-h light/12-h dark cycle. Mice had ad libitum access to standard laboratory chow (MFG; Oriental Yeast, Shiga, Japan; 3.57 kcal/g) and water unless otherwise specified.

Only male mice were used, as male-biased susceptibility to the social behavioral consequences of prenatal VPA exposure has been reported previously^13,43^.

### Method Details

#### VPA-treated mice

To generate VPA-treated mice^13,19^, pregnant females received a single intraperitoneal injection of sodium valproate (Sigma, #P4543) at 500 mg/kg on gestational day (GD) 12.5. Gestational timing was determined by the presence of a vaginal plug, designated as GD 0.5. Saline-treated control females received 10 mL/kg sterile saline at the same gestational stage.

### PLX5622- and minocycline-treated mice

PLX5622 (TargetMol, cat. #T7100; 90 mg/kg), minocycline (Fujifilm Wako, cat. #135-18671; 33 mg/kg), or tap water (control) was administered by oral gavage from four to six weeks of age. Mice receiving minocycline were used for experiments immediately at six weeks of age. In contrast, mice treated with PLX5622 or tap water underwent a one-week recovery period after the final administration before behavioral or histological analyses.

### Behavioral assays

The three-chamber test was conducted using an apparatus comprising three chambers (20 × 30 × 30 cm; width × length × height), with the side chambers connected to the central chamber by a 5 × 3 cm (width × height) passage. The test followed established protocols^44^. Animals were first habituated to the empty apparatus for 10 min. Grids were then placed in both side chambers, followed by an additional 10 min of habituation. A novel C57BL/6 young adult mouse or a Lego block was placed on a grid, and behavior was recorded for 10 min using a GoPro8 camera (GoPro; #CHDHX-801-FW). Experiments were conducted around zeitgeber time (ZT) 6, with ZT 0 defined as light onset. Social behavior was assessed by manual quantification of time spent exploring the grid containing the unfamiliar mouse versus the object. Investigation time was defined as active sniffing or exploration of the grid. Side chambers were cleaned with ethanol between sessions.

The marble-burying assay was performed following established protocols^35^ in a 14 × 32 × 14 cm (width × length × height) cage containing approximately 4 cm of wood-chip bedding. Animals were habituated to the empty cage for 10 min, after which 20 marbles were placed in the cage, and behavior was recorded for 10 min. At the end of the test, animals were removed, and buried marbles were counted manually. A marble was considered buried if at least two-thirds of its height was covered by bedding when viewed laterally.

### snRNAseq: data analysis

We reanalyzed the dataset reported previously^12^. Four batches were used: control, VPA-treated, VPA + CNO (PND 2) batch 1, and VPA + saline (PND 2). A second VPA + CNO (PND 2) batch was excluded because a substantial number of low-quality neurons were intermixed with non-neuronal clusters.

The data were imported into Seurat (R package v5.0.1), normalized using *NormalizeData* and *sctransform*, and integrated. Principal component analysis was performed using *RunPCA*, and clustering was conducted using the top 50 principal components at a resolution of 0.02 to achieve coarse separation of major cell types (Figure S1). For microglial analysis, *Ptprc*-positive clusters were extracted using *subset* function and re-clustered with the top 20 principal components at a resolution of 1.0. Only clusters expressing the microglial marker genes *Cx3cr1* and *P2ry12* were retained. For analyses shown in Figure 1, only control and VPA-treated batches were used, whereas Figure 3 analyses included only the VPA + CNO (PND 2) batch 1 and the VPA + saline (PND 2) batches.

### snRNAseq: DEGs, GO, and pathway analysis

A gene was considered expressed in a given cell type if at least one unique molecular identifier was detected in 30% or more of cells of that type. DEGs were identified as genes upregulated or downregulated under the specified comparison conditions and defined by an absolute log_2_ fold change (|log2-FC|) > 0.5 and p < 0.05. A curated list of high-confidence autism-associated genes was obtained from the SFARI Gene database (https://gene.sfari.org) as of 2 February 2025.

For Figures 1, 3, 4, and Figure S1, GO and pathway analyses of biological processes were performed for each cell cluster using *clusterProfiler*^21–23^ based on upregulated or downregulated DEGs in the corresponding comparisons. GO terms and pathways were derived from established curated terminologies. When a gene mapped to multiple isoform-specific pathways, these were consolidated and treated as a single pathway.

For Figure S1G–I, transcription factor activity analysis was performed using decoupleR (R package v2.9.7)^24^. For Figure S5, interactions between OT neurons and microglia were estimated using NeuronChat^31^ (R package v1.0.0).

### Histology and histochemistry

Brains from *OT-Cre*, *OT-Cre; Ai9*, *Oxtr-Cre*; *Ai9*, and *OT-Cre; stop-hM3Dq* mice were processed for immunolabeling. Mice were anesthetized with an overdose of isoflurane and perfused transcardially with PBS followed by 4% paraformaldehyde (PFA) in PBS. Brains were post-fixed overnight in 4% PFA at 4 °C, cryoprotected in 30% sucrose in PBS for at least 24 h, and embedded in O.C.T. compound (Tissue-Tek, cat#4583). Coronal sections (30 μm) spanning the brain were prepared using a cryostat (Leica, CM1860) and mounted on MAS-coated glass slides (Matsunami, cat#MAS-13). Primary antibodies used were rabbit anti-OT (1:500, ImmunoStar, cat#20068), goat anti-mCherry (1:500, Acris Antibodies GmbH, cat#ACR-AB0040-200-0.6), and chicken anti-IBA1 (1:500, Synaptic Systems GmbH, cat#234009). Secondary antibodies were donkey anti-rabbit Alexa Fluor 488 (1:250, Invitrogen, cat#A32790), donkey anti-goat Alexa Fluor 555 (1:500, Invitrogen, cat#A32816), goat anti-chicken Alexa Fluor 647 (1:500, Jackson ImmunoResearch, cat#103-605-155), and donkey anti-rabbit Alexa Fluor 647 (1:250, Invitrogen, cat#A31573). Sections were counterstained with DAPI (2.5 μg/mL) and imaged using an Olympus BX53 microscope with a 10× objective (numerical aperture 0.4).

### ISH

Fluorescent ISH was performed as described previously^45^. Mice were deeply anesthetized with isoflurane and perfused with PBS followed by 4% PFA. Brains were post-fixed overnight in 4% PFA, and 30-μm coronal sections were prepared using a cryostat (Leica). To generate cRNA probes, DNA templates were amplified by polymerase chain reaction from the C57BL/6J mouse genome or whole-brain cDNA (Genostaff, cat#MD-01). A T3 RNA polymerase recognition site (5ʹ-AATTAACCCTCACTAAAGGG) was added to the 3ʹ end of reverse primers. Primer sets for cRNA probe templates were as follows (forward primer first, reverse primer second):

*Il10ra*−1 5ʹ-CCAGGATGGGTTACTTGTGG; 5ʹ-TCTCAAAGAGCTTGCCCAGT

*Il10ra*−2 5ʹ-TCCCAAAGGTGTCACTAGAGC; 5ʹ-CTGCAGAAAAGGACCCACAT

*Il10ra*−3 5ʹ-GATGTTGTCGCGTTTGCTC; 5ʹ-CTCCAGGTCCACGATGTGA

*Nrxn1*−1 5ʹ-TGGTATATCTGGGAGGCTTGTT; 5ʹ-ACAGGTGTATTTAACTTTAATCCCATT

*Nrxn1*−2 5ʹ-TTACTCTGGCTGCTGCAATG; 5ʹ-CAATTCAATTTAGTTTTCCACCA

*Nrxn1*−3 5ʹ-TGAGAGAGAGGCAACGGTTT; 5ʹ-CGCAGTGATTTGTGTTGTGA

*Epha7*−1 5ʹ-TTCGTCTGAAGATGCTGGTG; 5ʹ-GGACGCAATCCATTTGAGTT

*Epha7*−2 5ʹ-GAAAACACCCCTGGGAACTT; 5ʹ-TGACACAGCCAGGATACCAA

*Epha7*−3 5ʹ-GCAGGCTATCAGCAAAAAGG; 5ʹ-CGAACACCATGAACACCAAG

DNA templates (500–1000 ng) were subjected to in vitro transcription using DIG RNA labeling mix (cat#11277073910) and T3 RNA polymerase (cat#11031163001) according to the manufacturer’s instructions (Roche Applied Science). When possible, up to three independent RNA probes were combined to enhance the signal-to-noise ratio.

For ISH combined with anti-mCherry staining, after hybridization and washing, sections were incubated overnight with horseradish peroxidase–conjugated anti-DIG (Roche, cat#11207733910; 1:500) and goat anti-mCherry (1:500, Acris Antibodies GmbH, cat#ACR-AB0040-200-0.6). Signals were amplified using TSA-plus Biotin (Akoya Biosciences, NEL749A001KT; 1:70 in 1× amplification diluent) for 25 min, washed, and mCherry-positive cells were visualized with donkey anti-goat Alexa Fluor 555 (1:500, Invitrogen, cat#A32816), whereas ISH probe-positive cells were detected with streptavidin–Alexa Fluor 488 (1:250; ThermoFisher Scientific, cat#S32354).

For ISH combined with anti-IBA1 staining, sections were incubated overnight with horseradish peroxidase–conjugated anti-DIG (Roche, cat#11207733910; 1:500) and chicken anti-IBA1 (1:500, Synaptic Systems GmbH, cat#234009). Signals were amplified using TSA-plus Biotin (Akoya Biosciences, NEL749A001KT; 1:70 in 1× amplification diluent) for 25 min, washed, and IBA1-positive cells were visualized with goat anti-chicken Alexa Fluor 647 (1:500, Jackson ImmunoResearch, cat#103-605-155), whereas ISH probe-positive cells were detected with streptavidin-Alexa Fluor 488 (1:250; ThermoFisher Scientific, cat#S32354).

### Xenium: sample preparation

Brains from saline- and VPA-treated *OT-Cre* mice were processed as follows. Mice were anesthetized with an overdose of isoflurane and perfused transcardially with PBS followed by 4% paraformaldehyde (PFA) in PBS. Brains were post-fixed overnight in 4% PFA at 4 °C and sequentially dehydrated in PBS, 70%, 80%, 90%, and 100% ethanol before dissection of the hypothalamic region containing the PVH. Tissues were infiltrated with paraffin and embedded. Prior to sectioning, paraffin blocks were rehydrated according to the Xenium protocol (CG000578) and sectioned at 5 μm. A subset of sections was stained with DAPI to identify the PVH, and adjacent sections were mounted onto Xenium slides. Xenium analysis was performed using the Prime 5K Mouse Pan Tissue and Pathways Assay Kit (cat#1000672) according to Xenium protocols CG000580, CG000760, and CG000584. In addition, 50 custom-designed genes were included to enable analyses of tissues beyond the scope of this project using the Xenium Add-on Custom 1 to 50 Gene Panel (cat#1000652); the full list of these genes is provided in Table S3.

### Xenium: data analysis

Xenium data were imported into Seurat (R package v5.1.0). Data from saline- and VPA-treated groups were normalized using *SCTransform* and integrated with *IntegrateLayers*. Clustering was performed using *RunPCA* with the top 50 principal components at a resolution of 1.8. Clusters were annotated based on characteristic marker genes (*Camk2a, Slc17a6, Slc32a1, Slc1a3, Sox10, Cx3cr1,* and *Foxj1*). Cluster identities generated in Seurat were exported as CSV files and imported into Xenium Explorer. Anterior and posterior PVH regions were defined based on DAPI signal and OT neuron distribution, and cell numbers within the PVH region of interest (ROI) were quantified per unit area. Cellular distributions within the ROI and expression patterns of selected RNAs were visualized using *ImageDimPlot* in Seurat.

### Quantification and statistics: snRNAseq and Xenium data

All statistical analyses of sequencing data were performed in R and RStudio. Analyses were conducted using RStudio Server 2023.06.1 Build 524 with R v4.3.1. GO term and pathway analyses were performed using RStudio 2022.12.0 + 353 with R v4.2.2. Statistical tests and criteria for snRNAseq analyses are described in the corresponding Methods sections. Numbers of biological replicates are provided in the respective figure legends.

### Quantification and statistics: histochemistry and ISH data

All image analyses were performed using *napari* (doi:10.5281/zenodo.3555620; version 0.4.15, Python 3.8.0, NumPy 1.23.1). To quantify anti-OT immunostaining and RNA probe signals in individual parvocellular PVH OT neurons, coronal sections 1000–1120 μm posterior to bregma were analyzed, based on prior work showing minimal presence of magnocellular PVH OT neurons in this region^12^. For magnocellular PVH OT neurons, coronal sections 520–640 μm posterior to bregma were analyzed. Using the *labels* tool in napari, all tdTomato-positive cells were selected as ROIs, and fluorescence intensity for each ROI was measured using *napari-skimage-regionprops* (version 0.5.3). Between 55 and 300 OT-positive ROIs were randomly collected from at least three animals per condition. Background fluorescence was subtracted, and corrected intensities were used for statistical analyses.

For RNA probe fluorescence in individual microglia, coronal sections 520–640 μm posterior to bregma were designated as anterior PVH, and sections 1000–1120 μm posterior to bregma as posterior PVH. The PVH was identified by DAPI staining and defined as the ROI. Fluorescence intensities of IBA1-positive cells within each ROI were quantified using *napari-skimage-regionprops* (version 0.5.3).

### Statistics

Statistical analyses were performed using Excel, Python, R, and js-STAR (v1.6.0). All tests were two-tailed unless stated otherwise. Sample sizes were not predetermined statistically but were chosen in accordance with standards in the field. Sample sizes and statistical tests are specified in the figure legends. Statistical significance was defined as p < 0.05 unless noted otherwise. Data are presented as mean ± standard error of the mean (SEM) unless indicated otherwise.

In box-and-whisker plots, the central line denotes the median, and the box boundaries represent the first and third quartiles. Whiskers extend to values within 1.5 times the interquartile range; values beyond this range were considered outliers and excluded from the plot. For experiments shown in Figures 2, 5, and Figures S1 and S4, data collection and analysis were performed by an experimenter blinded to experimental conditions.

## Data availability

Xenium data generated in this study have been deposited in the GEO repository (GSE318415). All data supporting the conclusions of this work are included in the main and supplementary figures.

## Code availability

This study does not report newly developed code. All Python and R scripts used for data analysis will be deposited elsewhere by publication.

## Acknowledgments

We thank members of the Miyamichi Lab for critical reading of the manuscript, the RIKEN BDR animal facility for animal care, and the RIKEN BDR sequencing facility for support with NGS and STx analyses. We also thank Yukiko U. Inoue and Takayoshi Inoue for sharing B6; C3-*Oxtr^em2(icre)Yinn^*/J mice, and Takeshi Sakurai for sharing *stop-hM3Dq* mice. This work was supported by the RIKEN Junior Research Associate Program (M.T.), JSPS KAKENHI (25K02368), JST CREST Program (JPMJCR2021), and the Takeda Science Foundation (K.M.).

## Author contributions

M.T. and K.M. conceived the study. M.T. performed the experiments and analyzed the data, with support from H.S. and M.H. D.C. and T.K. assisted with 10X Xenium operation. M.T. and K.M. wrote the manuscript with input from all authors.

## Supplemental Information

**Figure S1:**
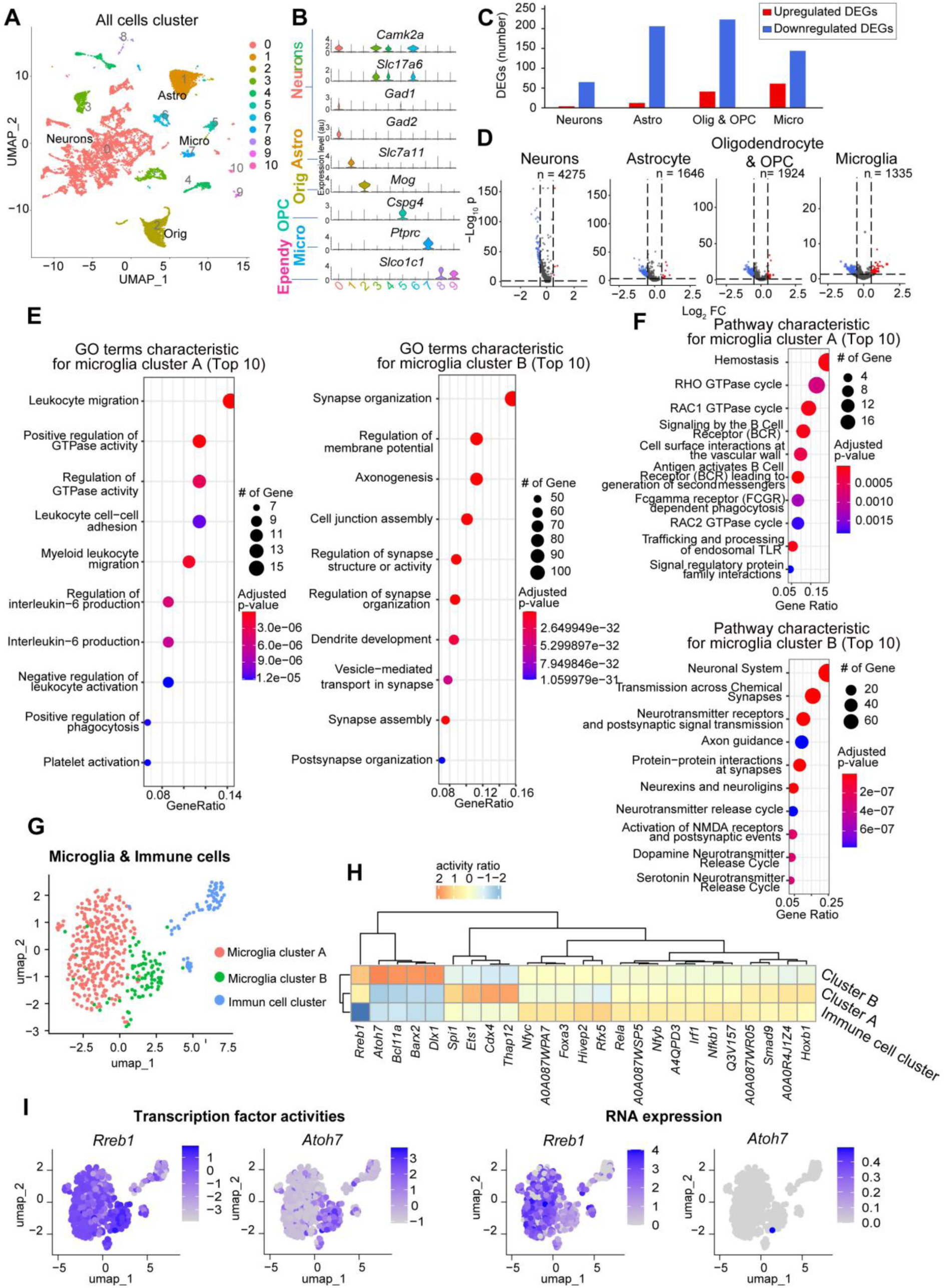
Impacts of embryonic VPA treatment on non-neuronal cells and classification of microglial subtypes, related to Figure 1. (A) UMAP representation of all cells analyzed, showing 10 clusters. (B) Violin plots of representative marker genes for major cell types. Based on these expression patterns, clusters 0, 3, 4, and 6 were annotated as neurons; cluster 1 as astrocytes; cluster 2 as oligodendrocytes; cluster 5 as OPCs; cluster 7 as microglia; and clusters 8 and 9 as ependymal cells. (C) Numbers of upregulated (red) and downregulated (blue) DEGs in neurons, astrocytes, oligodendrocytes/OPCs, and microglia. (D) Volcano plots showing DEGs (upregulated in the VPA-treated group, red; downregulated in the VPA-treated group, blue) in neurons, astrocytes, oligodendrocytes/OPCs, and microglia, as in Figure 1D. P-values were calculated using the Wilcoxon rank-sum test without correction for multiple comparisons. (E) Heatmap showing p-values for the top 10 GO terms enriched among characteristic genes of each microglial cluster. P-values were calculated using one-sided Fisher’s exact test with Benjamini–Hochberg correction. (F) Heatmap showing p-values for the top 10 enriched pathways identified from representative genes of each microglial cluster. P-values were calculated using one-sided Fisher’s exact test with Benjamini–Hochberg correction. (G) UMAP representation of microglia and immune cell clusters. (H) Heatmaps of transcription factor activities in microglial clusters A and B and the immune cell cluster. (I) Transcription factor activities (left) and RNA expression (right), with color scales indicating log-normalized expression.

**Figure S2:**
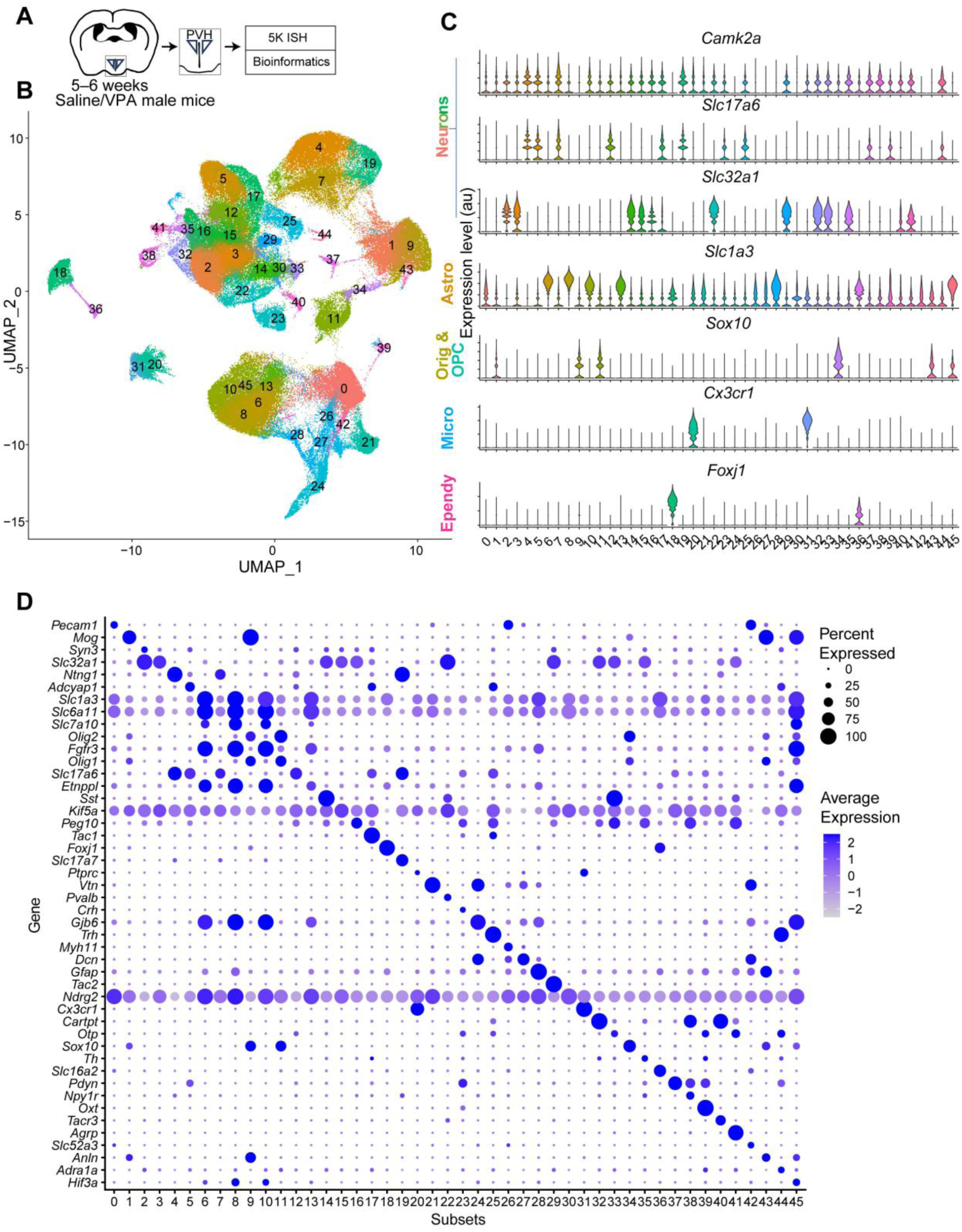
Global classification of Xenium spatial transcriptomics data, related to Figure 2. (A) Schematic of the Xenium data collection and analysis workflow. (B) UMAP representation of all profiled cells, revealing 46 transcriptomic subtypes. (C) Violin plots of canonical marker genes used to classify major cell types. Based on marker expression, subtypes 2, 3, 4, 5, 7, 12, 14, 15, 16, 17, 19, 22, 23, 25, 29, 30, 32, 33, 35, 37, 38, 39, 40, 41, and 44 were annotated as neurons; subtypes 0, 6, 8, 10, 13, 21, 24, 26, 27, 28, 42, and 45 as astrocytes (Astro); subtypes 1, 9, 34, and 43 as oligodendrocytes (Olig); subtype 11 as OPCs; subtypes 20 and 31 as microglia (Micro); and subtypes 18 and 36 as ependymal cells (Ependy). (D) Enriched differentially expressed genes across all subtypes. Dot size reflects population size, and the heatmap indicates log-normalized expression.

**Figure S3:**
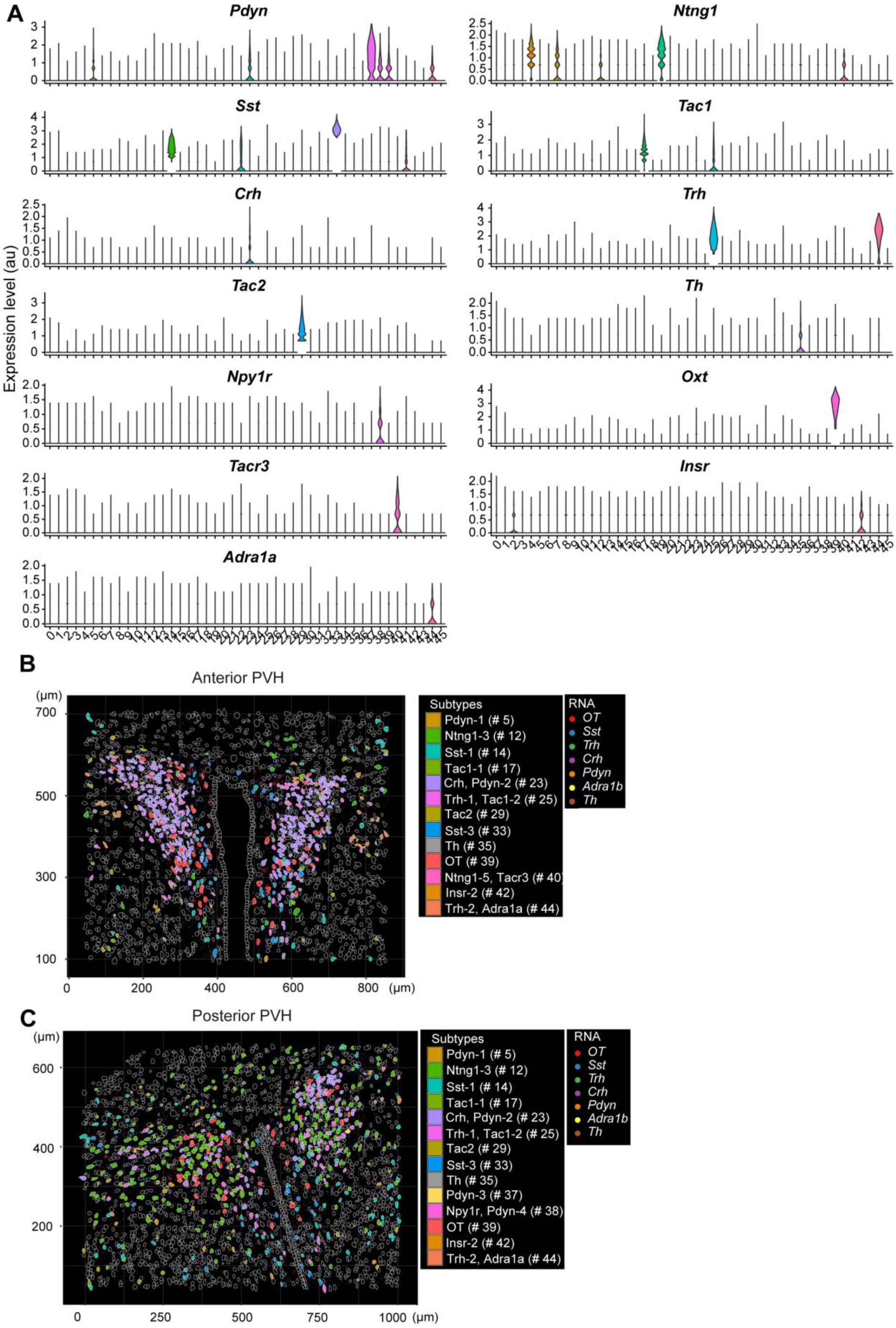
Spatial distribution of PVH neuronal subtypes identified by Xenium, related to Figure 2. (A) Violin plots showing expression of canonical marker genes for major PVH neuronal subtypes. Based on marker expression, subtypes 5, 23, 37, 38, 39, and 44 were annotated as *Pdyn* neurons; subtypes 14, 22, 33, and 41 as *Sst* neurons; subtype 23 as *Crh* neurons; subtype 29 as *Tac2* neurons; subtype 38 as *Npy1r* neurons; subtype 40 as *Tacr3* neurons; subtype 44 as *Adra1a* neurons; subtypes 4, 7, 12, 18, and 40 as *Ntng1* neurons; subtypes 17 and 25 as *Tac1* neurons; subtypes 25 and 44 as *Trh* neurons; subtype 35 as *Th* neurons; subtype 39 as *OT* neurons; and subtype 42 as *Insr* neurons. With the probe set used, magnocellular and parvocellular OT neurons could not be distinguished. (B, C) Spatial maps of major PVH neuronal subtypes in the anterior (B) and posterior (C) PVH. Colors denote 14 neuronal subtypes; dots represent RNA signals (red, *OT*; indigo, *Sst*; green, *Trh*; purple, *Crh*; orange, *Pdyn*; yellow, *Adra1b*; brown, *Th*). STx data further revealed that *Sst* and *Trh* neurons each comprised two spatially distinct subtypes, one located closer to the third ventricle and another positioned more laterally (B). Previous snRNAseq studies indicate that these neuronal populations comprise multiple transcriptomic subtypes^12,46^, consistent with the present STx observations.

**Figure S4:**
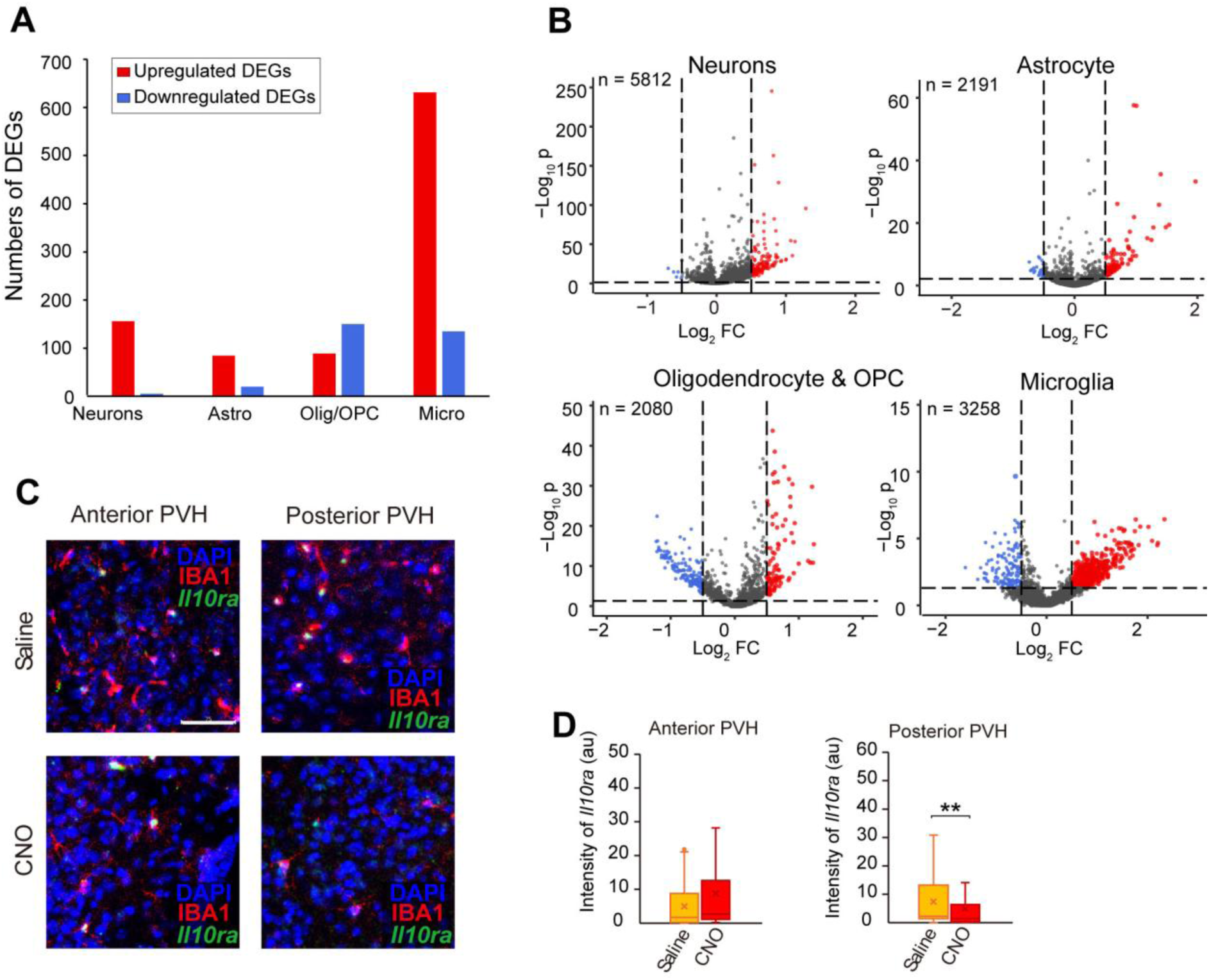
Impact of neonatal OT-neuron stimulation on non-neuronal PVH cell types and additional histochemical assays, related to Figure 3. (A) Numbers of upregulated (red) and downregulated (blue) DEGs identified in neurons, astrocytes, oligodendrocytes/OPCs, and microglia. (B) Volcano plots showing DEGs (upregulated in the CNO-treated group, red; downregulated in the CNO-treated group, blue) in neuronal, astrocytic, oligodendrocyte/OPC, and microglial clusters, as in Figure 3D. P-values were calculated using the Wilcoxon rank-sum test without adjustment for multiple comparisons. (C) Representative coronal sections from saline- and VPA-treated mice at anterior and posterior PVH levels, showing anti-IBA1 immunostaining and *Il10ra* expression detected by ISH. Scale bar, 75 μm. (D) Fluorescence intensities of *Il10ra* ISH signals. A total of 47–52 cells from three animals per condition were analyzed. p < 0.01 by the two-sided Wilcoxon rank-sum test. Box plots follow the definitions in Figure 2F.

**Figure S5:**
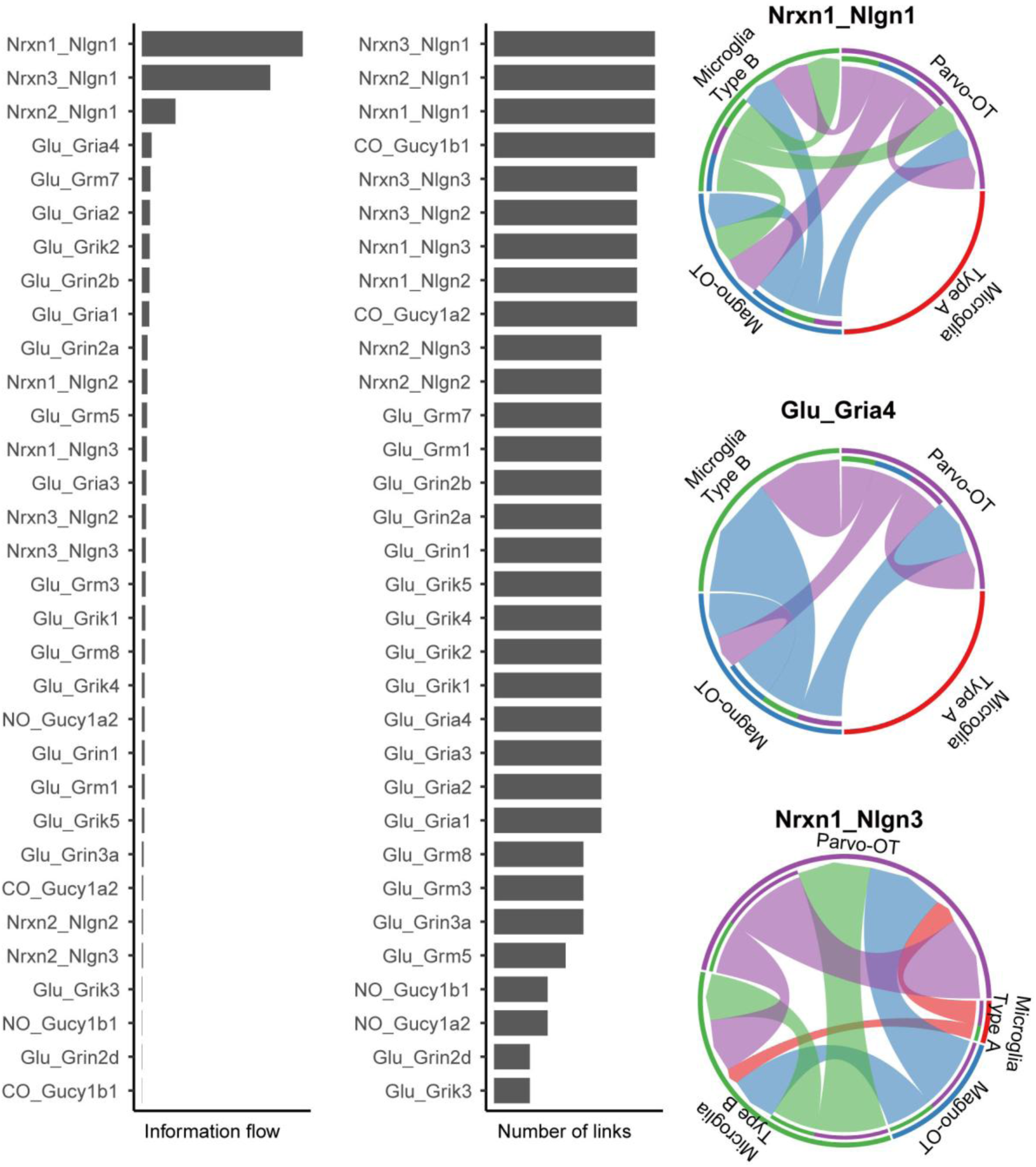
Estimated cell–cell communication inferred from snRNAseq data, related to Figure 4. Left: Bar plots showing information flow and numbers of predicted ligand–receptor links among parvocellular (Parvo) OT neurons, magnocellular (Magno) OT neurons, microglial cluster A, and microglial cluster B, as inferred using NeuronChat^31^. Right: Circular chord diagrams illustrating representative ligand–receptor interactions, including *Nrxn1-Nlgn1*, *Glu-Gria4*, and *Nrxn1-Nlgn3* pairs.

**Figure S6:**
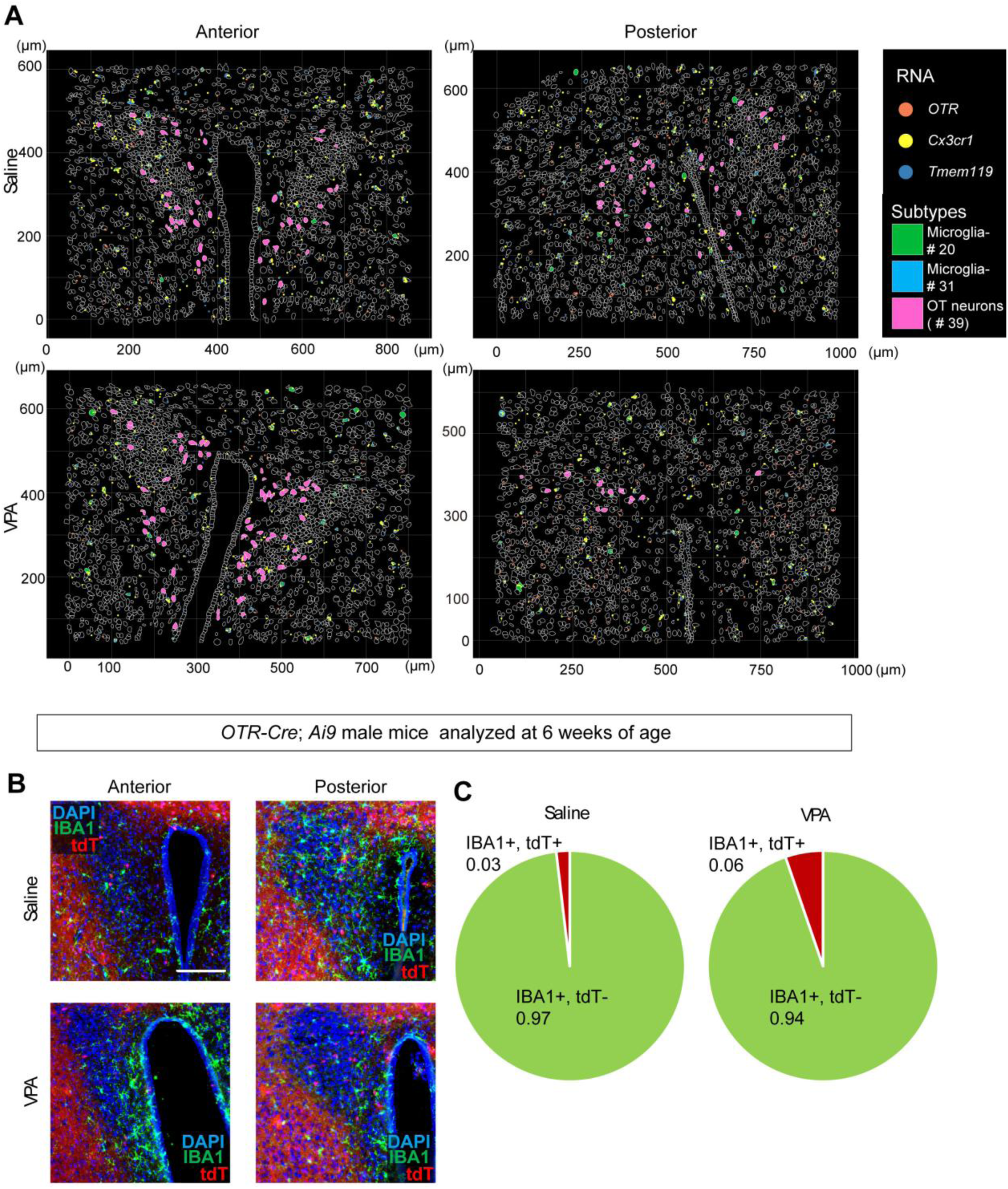
Spatial distribution of microglia and *OTR*-positive cells, related to Figure 4. (A) Spatial map of OT neurons (pink) and two microglial subtypes (green and blue) identified using the 10X Xenium platform. White contours delineate detected cells in the anterior and posterior PVH. Dots indicate individual RNA signals (orange, *OTR*; yellow, *Cx3cr1*; indigo, *Tmem119*). (B) Representative coronal PVH sections from male *Otr-Cre*; *Ai9* mice showing tdTomato-positive cells (tdT, red) and anti-IBA1 immunostaining (green), counterstained with DAPI. Scale bar, 200 μm. (C) Pie charts showing proportions of IBA1-positive/tdT-positive and IBA1-positive/tdT-negative cells in saline- and VPA-treated mice. These data do not support widespread OTR expression in PVH microglia.

**Figure S7:**
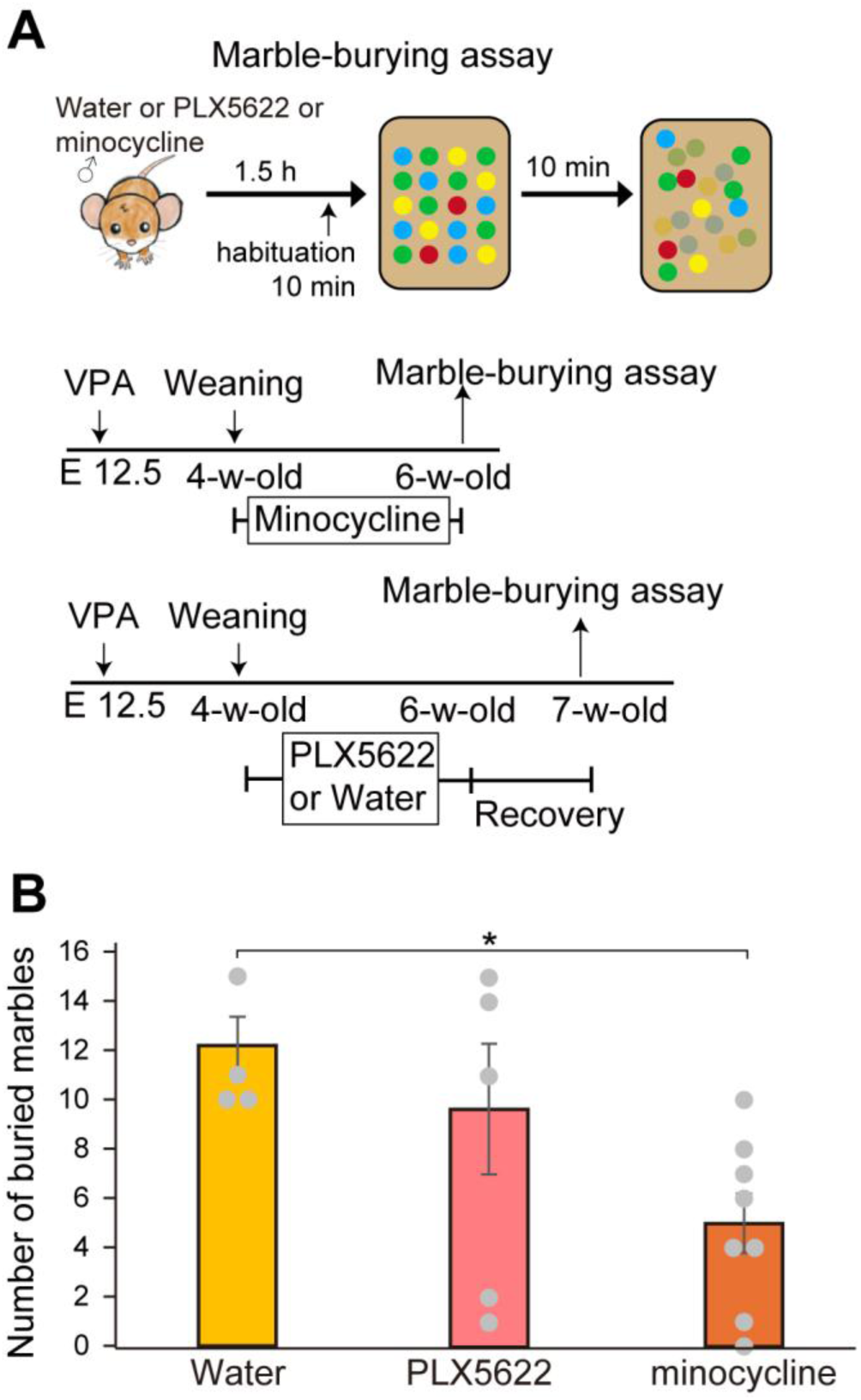
Impact of microglial manipulation on marble-burying behavior, related to Figure 5. (A) Top: Schematic of the marble-burying assay. Bottom: Experimental timeline. (B) Number of buried marbles across the three experimental groups. *p* < 0.05 by the Wilcoxon rank-sum test with Bonferroni correction.

**Figure S8:**
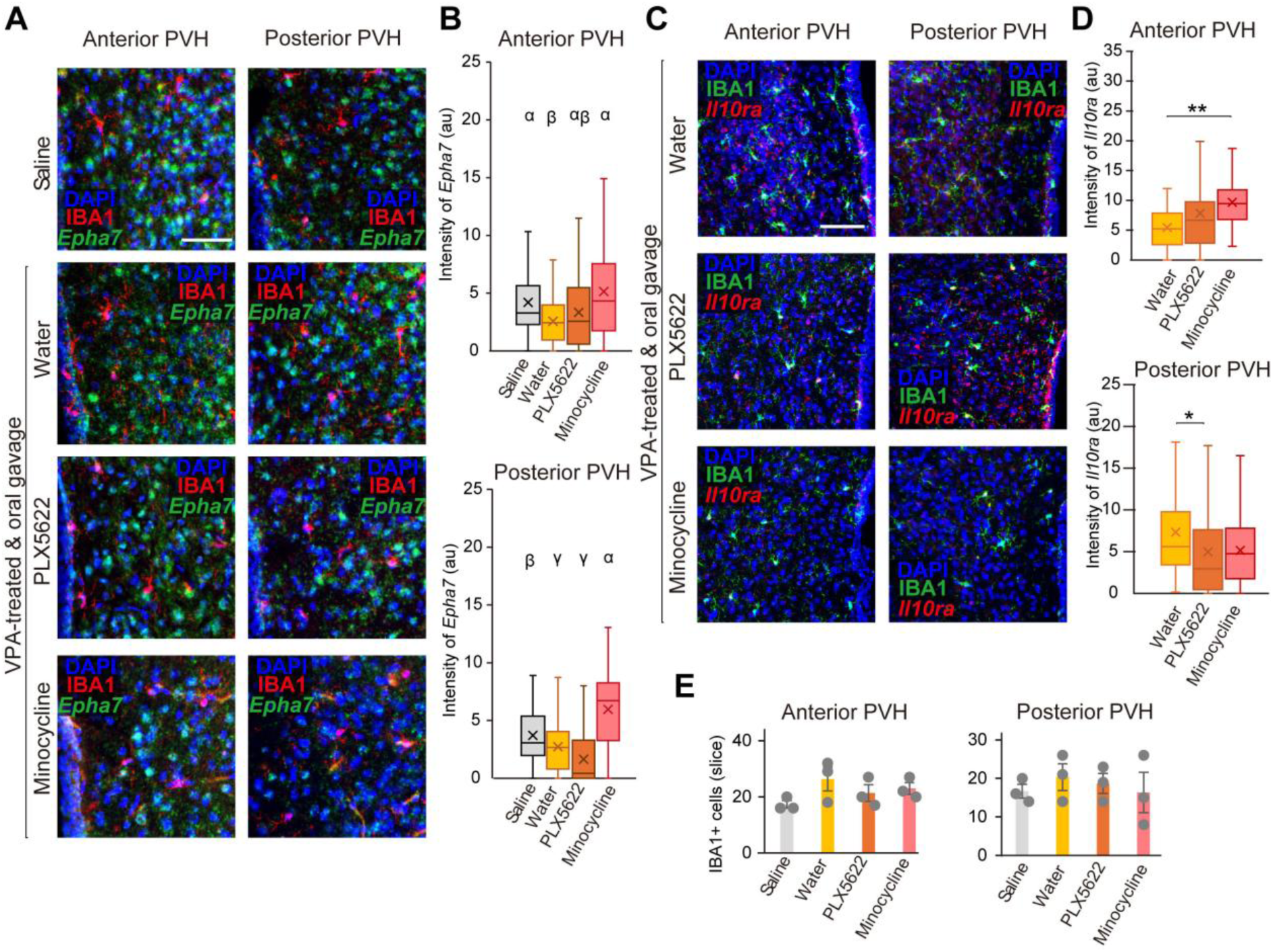
Impact of microglial manipulation on *Epha7* and *Il10ra* expression in PVH microglia, related to Figure 5. (A) Representative coronal anterior and posterior PVH sections from embryonic saline-treated controls (top), VPA- and water-treated mice (second row), VPA- and PLX5622-treated mice (third row), and VPA- and minocycline-treated mice (bottom). Sections show anti-IBA1 immunostaining and *Epha7* expression detected by ISH, counterstained with DAPI. Scale bar, 75 μm. (B) Quantification of *Epha7* ISH signal intensity across conditions. A total of 49–79 microglial cells from N = 3 mice were analyzed. Different letters (α, β, γ) indicate p < 0.05 by the two-sided Wilcoxon rank-sum test with Bonferroni correction. (C) Representative anterior and posterior PVH sections from microglial manipulation conditions showing anti-IBA1 immunostaining and *Il10ra* expression detected by ISH, counterstained with DAPI. Scale bar, 100 μm. (D) Quantification of *Il10ra* ISH signal intensity across conditions. A total of 53–97 microglial cells from N = 3 mice were analyzed. p < 0.05, p < 0.01 by the Wilcoxon rank-sum test with Bonferroni correction. Box plots follow the definitions in Figure 2F. (E) Numbers of IBA1-positive cells in the anterior and posterior PVH across treatment conditions. N = 3 animals per group.

**Figure S9:**
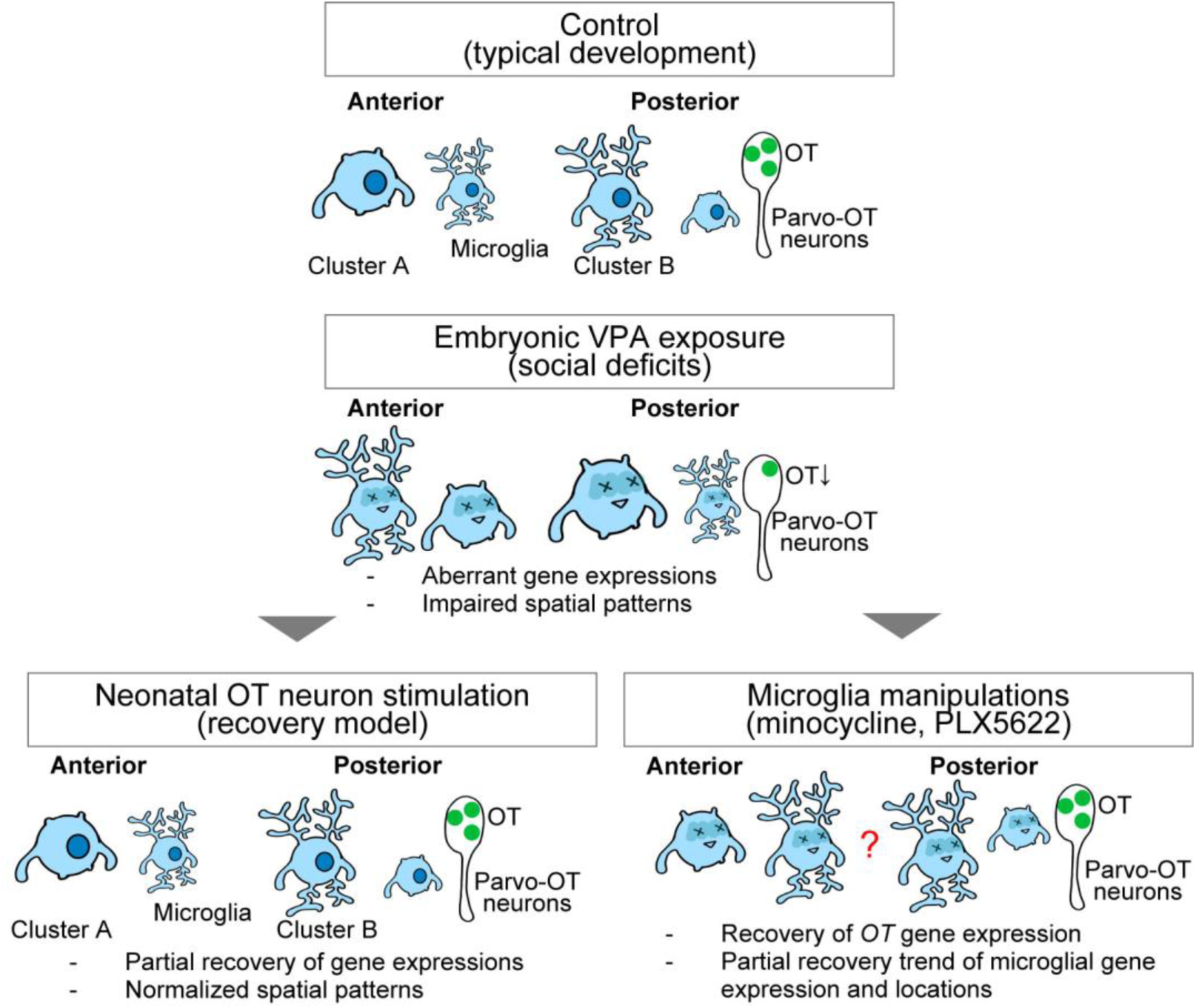
Graphical abstract, related to Figures 1–5. snRNAseq and spatial transcriptomics identified two PVH microglial subtypes: cluster A, an immune-like population, and cluster B, a neuron-associated population. These subtypes exhibited spatial segregation, with cluster A enriched in the anterior PVH and cluster B preferentially localized to the posterior PVH. Embryonic VPA exposure disrupted this organization and broadly downregulated gene expression in both clusters. Neonatal OT-neuron stimulation partially restored aberrant microglial transcriptional states and spatial distribution. In addition, pharmacological microglial manipulation increased OT expression in putative parvocellular OT neurons and partially normalized microglial positioning and gene expression.

